# ER transmembrane protein TMTC3 contributes to O-mannosylation of E-cadherin, Cellular Adherence and Embryonic Gastrulation

**DOI:** 10.1101/822270

**Authors:** Jill B. Graham, Johan C. Sunryd, Ketan Mathavan, Emma Weir, Ida Signe Bohse Larsen, Adnan Halim, Henrik Clausen, Hélène Cousin, Dominque Alfandari, Daniel N. Hebert

**Affiliations:** Department of Biochemistry and Molecular Biology, University of Massachusetts, Amherst, Amherst, MA USA; Department of Veterinary and Animal Sciences, University of Massachusetts, Amherst, Amherst, MA USA; Program in Molecular and Cellular Biology, University of Massachusetts, Amherst, Amherst, MA USA; Department of Cellular and Molecular Medicine, University of Copenhagen, Copenhagen, Denmark; Copenhagen Center for Glycomics, University of Copenhagen, Copenhagen, Denmark

**Author notes:** To whom correspondence should be addressed: Daniel N. Hebert, Department of Biochemistry and Molecular Biology, University of Massachusetts, 240 Thatcher Road Amherst, MA 01003.

**Keywords:** endoplasmic reticulum, O-glycosylation, O-mannosylation, cell adhesion, Cobblestone lissencephaly, protein modification

## Abstract

Protein glycosylation plays essential roles in protein structure, stability and activity such as cell adhesion. The cadherin superfamily of adhesion molecules carry O-linked mannose glycans at conserved sites and it was recently demonstrated that the *TMTC1-4* genes contribute to the addition of these O-linked mannoses. Here, biochemical, cell biological and organismal analysis was used to determine that TMTC3 supports the O-mannosylation of E-cadherin, cellular adhesion and embryonic gastrulation. Using genetically engineered cells lacking all four *TMTC* genes, overexpression of TMTC3 rescued O-linked glycosylation of E-cadherin and cell adherence. The knockdown of the Tmtcs in *Xenopus laevis* embryos caused a delay in gastrulation that was rescued by the addition of human TMTC3. Mutations in *TMTC3* have been linked to neuronal cell migration diseases including Cobblestone lissencephaly. Analysis of TMTC3 mutations associated with Cobblestone lissencephaly found that three of the variants exhibit reduced stability and missence mutations were unable to complement TMTC3 rescue of gastrulation in *Xenopus* embryo development. Our study demonstrates that TMTC3 regulates O-linked glycosylation and cadherin-mediated adherence, providing insight into its effect on cellular adherence and migration, as well the basis of TMTC3-associated Cobblestone lissencephaly.

## Introduction

Protein glycosylation is the most common and diverse co/post-translational protein modification (Freeze and Elbein, 2009). Carbohydrates play general metabolic, structural and biophysical roles in the cell (O’Connor and Imperiali, 1996; Apweiler *et al.*, 1999; Solá and Griebenow, 2009; Varki and Sharon, 2009; Price *et al.*, 2012; Hebert *et al.*, 2014). There are two general types of glycosylation that are categorized by the type of bond used to covalently attach the carbohydrate to the protein (Imperiali and Hendrickson, 1995; Jayaprakash and Surolia, 2017). N- and O-linked glycans are attached to proteins through N-glycosidic (Asn) or O-glycosidic (Ser/Thr) bonds, respectively. A third of the proteome is targeted to the secretory pathway in eukaryotic cells where both N- and O-linked modifications are commonly observed (Wallin and von Heijne, 1998; Chen *et al.*, 2005; Choi *et al.*, 2010). The endoplasmic reticulum (ER) serves as the entry portal for secretory pathway cargo. The majority of the proteins in the ER are modified by N-linked glycans; however, a restricted fraction is modified by O-linked glycans (Apweiler *et al.*, 1999; Vester-Christensen *et al.*, 2013).

A diverse set of O-glycans are added in the ER that include mannose, fucose, glucose or N-acetyl glucosamine (Joshi *et al.*, 2018). O-mannosylation is conserved from fungi to mammals. It is initiated by the transfer of a mannose saccharide from a donor dolichol P-mannose substrate embedded in the ER membrane to a Ser/Thr on the target protein (Bause and Lehle, 1979). This transfer is catalyzed by ER resident protein O-mannosyltransferases most commonly involving POMT1 and POMT2 with α-dystroglycan as the main substrate in mammals, and Pmt1-7 in yeast (Wing *et al.*, 1992; Strahl-Bolsinger *et al.*, 1993; Immervoll *et al.*, 1995; Lussier *et al.*, 1995; Guerreiro *et al.*, 1996). Their general architecture is that of large polytopic membrane proteins, which display variations of a diacidic motif (i.e. Asp-Glu) in the first luminal loop region proximal to the membrane (Liu and Mushegian, 2003; Lairson *et al.*, 2008; Loibl and Strahl, 2013; Bai *et al.*, 2019)(Supplemental Fig. 1B). Metazoan and yeast O-mannosyltransferases also contain conserved <u>m</u>annosyltransferase, inositol triphosphate and ryanodine receptor (MIR) domains. MIR domains have been shown to be essential for enzymatic activity in yeast and have been implicated in substrate binding based on studies performed on other mammalian MIR domain containing proteins such as SDF2 (Girrbach *et al.*, 2000; Ponting, 2000; Meunier *et al.*, 2002; Lommel *et al.*, 2011; Fujimori *et al.*, 2017).

A new putative family of O-mannosyltransferases was recently discovered in mammals (Larsen *et al.*, 2017a, 2017b). This family is comprised of four tetratricopeptide repeat (TPR)-containing proteins that appear to be ER polytopic transmembrane proteins called TMTC1-4 (trans<u>m</u>embrane and <u>T</u>PR-<u>c</u>ontaining proteins 1-4)(Della-Morte *et al.*, 2011; Racapé *et al.*, 2011; Cao *et al.*, 2012; Sunryd *et al.*, 2014; Larsen *et al.*, 2017a; Li *et al.*, 2018). TMTC1, 2 and 4 are ER proteins involved in Ca^2+^ regulation as they associate with ER Ca^2+^ re-uptake pump SERCA2B (Sunryd *et al.*, 2014; Li *et al.*, 2018). The knockouts of *TMTC1-4* have been shown using glycoproteomics to be involved in the O-mannosylation of cadherins (Larsen *et al.*, 2017a, 2019). TMTC1-4 contain homologous structural elements as their N-terminal halves are comprised of a number of hydrophobic domains and the C-terminal halves are predicted to be composed mainly of long stretches of TPR motifs (8-12) (Karpenahalli *et al.*, 2007; Letunic and Bork, 2018)(Fig. 1A). Two of the TMTCs, TMTC3 and TMTC4, contain putative diacidic motifs (Asp-Asp or Glu-Glu) on N-terminal luminal loops based on predictive topology, making them appear homologous to the POMTs and Pmts (Hessa *et al.*, 2007; Bai *et al.*, 2019)(Supplemental Fig. 1B and C).

**Figure 1.**
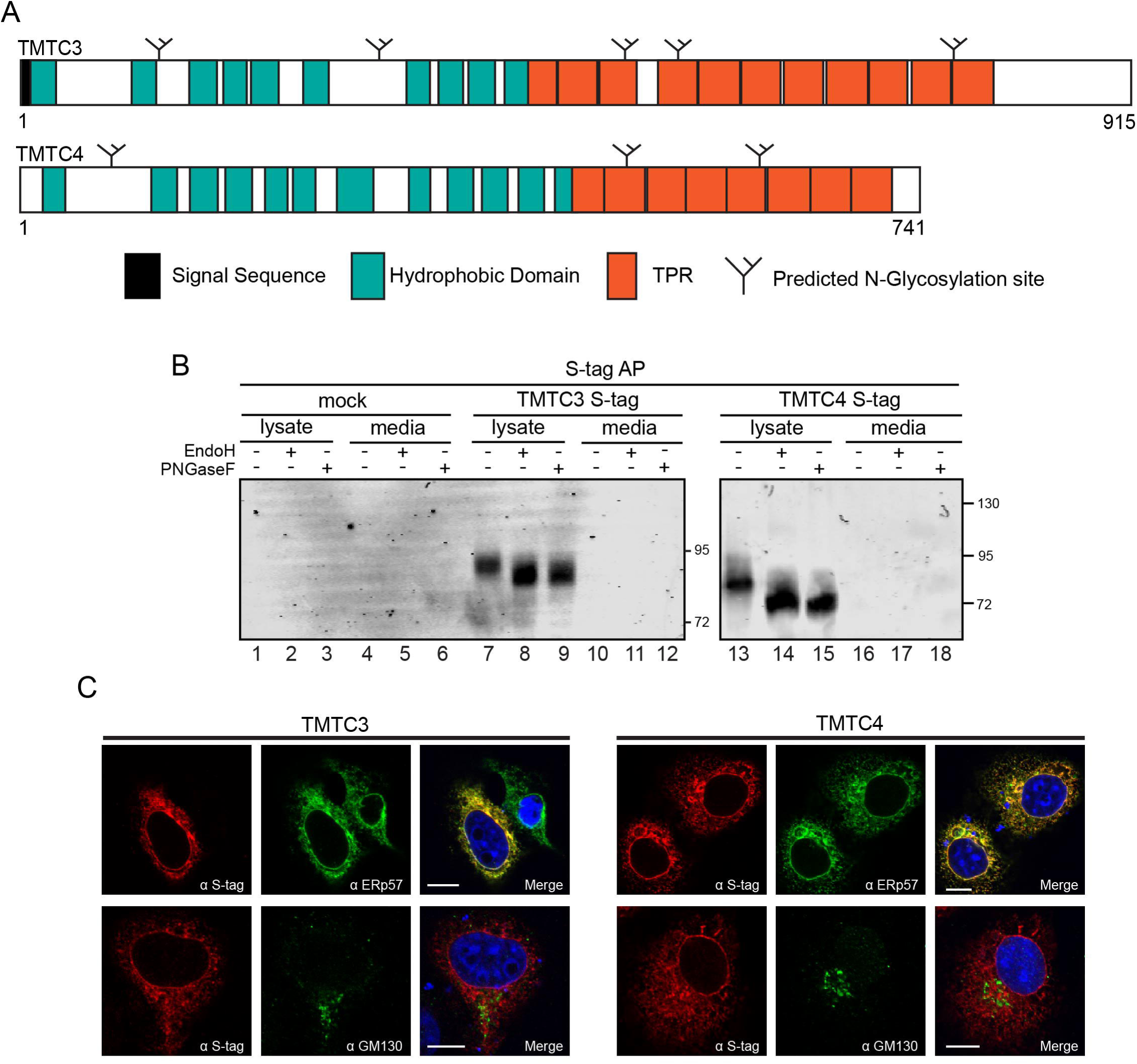
TMTC3 and TMTC4 are ER resident proteins. (A) The organization of TMTC3 and TMTC4 with signal sequences (black), hydrophobic domains (teal) and TPR domains (orange) as designated. The start and end positions of each domain are as follows: TMTC3 hydrophobic domains (aa 9-30, 94-115, 140-161, 169-187, 192-214, 235-257, 321-340, 347-367, 371-393 and 402-422), TMTC3 TPR motifs (aa 412-445, 446-479, 480-513, 529-562, 563-596, 597-630, 632-665, 669-702, 703-736, 738-771 and 772-805), TMTC4 hydrophobic domains (aa 20-38, 109-130, 142-161, 170-192, 203-221, 225-245, 269-291, 320-339, 353-374, 381-403, 412-433 and 440-460) and TMTC4 TPR motifs (aa 448-481, 482-515, 516-549, 550-583, 584-617, 618-651, 652-685 and 686-719). Predicted endogenous N-linked glycosylation sites are indicated by small black, branched structures. (B) HEK-293T cells were transfected with S-tagged TMTC3 or TMTC4 as indicated and were affinity purified from the cell lysate and media using S-protein agarose. Samples were then subjected to a glycosylation assay with either EndoH (lanes 2, 5, 8, 11, 14 and 17) or PNGaseF (lanes 3, 6, 9, 12, 15 and 18) digestion as indicated. Reducing sample buffer was added and the samples were analyzed by a 6% (TMTC3 S-tag) and 8% (TMTC4 S-tag) SOS-PAGE. (C) Cellular localization of TMTC3 and TMTC4 was investigated by confocal microscopy. COS7 cells were transfected with TMTC3 or TMTC4 cDNA. Fixed cells were stained with S-tag, ERp57 (ER) or GM130 (Golgi) antisera. Nuclei were visualized by DAPI staining (blue). Scale bars correspond to 10 μm.

TPRs are protein structural motifs that support protein-protein interactions and are frequently found on co-chaperones (Magliery and Regan, 2005; Brehme *et al.*, 2014). A single TPR motif consists of a degenerate 34-amino acid sequence that is comprised of two anti-parallel α-helices (D’Andrea and Regan, 2003; Magliery and Regan, 2005). TPR domains are most commonly found in a cluster of three TPR motifs; however, clusters comprised of up to sixteen sequential TPRs have been observed (D’Andrea and Regan, 2003). Proteins with three TPRs in a cluster favor the recognition of short and defined sequences, whereas proteins with long stretches of consecutive TPRs tend to be more promiscuous in their selectivity. Unlike classic TPR co-chaperones, which contain three to four consecutive motifs, the composition of TMTC1-4, with eight to twelve consecutive motifs, is similar to that of the well-studied TPR-containing cytoplasmic O-GlcNAc transferase (OGT), which contains a stretch of 12.5 TPRs followed by a C-terminal catalytic domain (Jínek *et al.*, 2004; Lazarus *et al.*, 2011). The structure of OGT reveals that the TPRs form a super helix responsible for homodimerization of OGT and are proposed to scaffold interactions with other proteins, potentially playing a role in substrate selectivity. Dimerization of the TPR domain is mediated by the convex faces of the superhelical monomers, and the concave surface contains conserved Asn, which form a continuous ladder. This Asn conservation is similarly observed in the ARM-repeat proteins importin-α and β-catenin (Conti *et al.*, 1998; Huber and Weis, 2001). The Asn in these proteins contribute to binding of a target peptide (nuclear localization sequence with importin-α and E-cadherin with β – catenin), suggesting that the TMTCs may use a similar mechanism of protein-protein interaction and substrate selection.

Mutations in the *TMTC* genes have been linked to various human disease states (Jerber *et al.*, 2016; Runge *et al.*, 2016; Farhan *et al.*, 2017; Li *et al.*, 2018). Point mutations in human *TMTC2* and the knockout of *Tmtc4* in mice result in hearing loss (Runge *et al.*, 2016; Guillen□Ahlers *et al.*, 2018; Li *et al.*, 2018). Ten mutations in *TMTC3* are associated with neuronal cell migration diseases (Jerber *et al.*, 2016; Farhan *et al.*, 2017). Homozygous and compound heterozygous mutations resulting in single amino acid or frame shift changes in TMTC3 were identified in a cohort of families with recessive forms of Cobblestone lissencephaly, a severe brain malformation in which over migration of neurons and glial cells results in the formation of cortical dysplasia or brain development abnormalities (Jerber *et al.*, 2016). Another study identified heterozygous variants of *TMTC3* in patients with periventricular nodular heterotopia (PVNH), a common brain malformation caused by the failure of neurons to migrate from the ventricular zone to the cortex (Farhan *et al.*, 2017). While mutation or loss of the *TMTC* genes has been associated with a number of diseases, an understanding of how these mutations result in specific defects is unclear.

Here, *in silico*, biochemical, cell and developmental biological approaches were used to expand our understanding of the organization, localization, activity and function of TMTC3 and TMTC4. Previously uncharacterized TMTC3 and 4 were identified as ER TPR-containing membrane proteins with their TPR domains orientated within the ER lumen. Using *TMTC* HEK293 knockout cells, it was demonstrated that TMTC3 complementation recovered the O-mannosylation of E-cadherin. While the knockout of the *TMTC*s did not affect the stability, localization or trafficking of E-cadherin, it did affect cellular adherence and the overexpression of TMTC3 was able to partially recover adherence, specifically E-cadherin mediated adherence. Subsequent analysis of protein O-mannosyltransferase protein architecture showed that TMTC3 shares functional similarity with O-mannosyltransferases (POMT1/2 and Pmt1-7) as it contains two diacidic motifs (Asp-Asp and Glu-Glu) located in a putative N-terminal luminal loop and mutation of the Asp-Asp motif results in reduced TMTC3 stability. Additionally, the knockdown of Tmtc3 and Tmtc4 in *Xenopus laevis* resulted in an embryonic gastrulation delay phenotype and the delay was rescued by human TMTC3. There are eight disease variants of TMTC3 recently associated with Cobblestone lissencephaly and two associated with PVNH (Jerber *et al.*, 2016; Farhan *et al.*, 2017). While novel biochemical characterization of eight of these disease variants showed that they all localize to the ER, three of them are less stable than wild type (WT) and none of the missense mutations are able to rescue the delay in gastrulation caused by the knockdown of Tmtc3 in *Xenopus laevis*. The identification of TMTC3’s role in O-glycosylation of E-cadherin in combination with its knockdown in *Xenopus* embryos provides further insight into the role O-glycosylation plays in cell-cell adhesion and migration, and the etiology of Cobblestone lissencephaly caused by *TMTC3* mutation.

## Results

### TMTC3 and TMTC4 are ER resident proteins

*In silico* analysis, using SignalP4.0, TargetP1.1, ΔG, TPRPred and domain architecture database SMART7, indicated that TMTC3 (NCBI Accession # <u>NP_861448.2.</u> (http://www.ncbi.nlm.nih.gov/protein/NP_861448.2)) and TMTC4 (NCBI Accession # <u>NP_001073137.1</u> (http://www.ncbi.nlm.nih.gov/protein/NP_001073137)) contained a potential N-terminal signal sequence, ten and twelve hydrophobic segments and eleven and eight C-terminal TPR motifs, respectively (Fig. 1A)(Nielsen *et al.*, 1997; Emanuelsson *et al.*, 2000; Hessa *et al.*, 2007; Karpenahalli *et al.*, 2007; Nielsen, 2017; Letunic and Bork, 2018; UniProt: a worldwide hub of protein knowledge, 2019). *TMTC3* and *TMTC4* cDNAs were subcloned into mammalian expression vectors encoding a C-terminal S-tag, and their cellular localization was determined by glycosylation assay and confocal immunofluorescence microscopy. Secretory proteins are commonly modified in the ER with N-linked glycans at the consensus site Asn-Xxx-Ser/Thr. TMTC3 and TMTC4 possess five and three predicted N-linked glycosylation consensus sites, respectively (Fig. 1A), therefore a glycosylation assay was used to further analyze ER targeting and localization (Gupta, 2002). As the molecular weight of an N-linked glycan is ∼2.5 kDa, the removal of N-linked glycans by glycosidase treatment results in a corresponding increase in mobility for the deglycosylated protein. Endoglycosidase H (Endo H) trims the high mannose glycans encountered in the ER while Peptide-N-Glycosidase F (PNGase F) removes complex glycans acquired in the Golgi in addition to high mannose glycans.

HEK293T cells were transfected with TMTC3 or TMTC4 containing C-terminal S-tags. Cell lysate and media fractions were affinity precipitated with S-protein agarose beads followed by glycosidase treatment. Shifts upon PNGaseF treatment (Fig. 1B, lanes 9 and 15) were observed for both TMTC3 and TMTC4 demonstrating that both proteins were targeted to the ER and received N-linked glycans. A similar increase in mobility was observed upon Endo H treatment (Fig. 1B, lanes 8 and 14) indicating that the carbohydrates were high mannose glycoforms suggesting that TMTC3 and TMTC4 are ER resident proteins (Fig. 1B).

COS7 cells were transfected with either TMTC3 S-tag or TMTC4 S-tag as COS7 cells are highly amenable to imaging. Fluorescence staining of TMTC3 and TMTC4 was compared against an ER (ERp57) or Golgi (GM130) marker (Fig. 1C). Both TMTC3 and TMTC4 co-localized with ERp57, while co-localization was not observed with GM130 (Fig. 1C). Therefore, the glycosylation profiles for TMTC3 and TMTC4 and the cellular distribution are consistent with TMTC3 and TMTC4 residing in the ER.

### TMTC3 and TMTC4 are ER membrane proteins with luminal orientated TPR motifs

Analysis of the TMTC3 and TMTC4 protein sequences with ΔG prediction demonstrated that they contained ten and twelve hydrophobic segments, respectively, that could potentially serve as transmembrane domains to create polytopic membrane proteins (Fig. 1A) (Hessa *et al.*, 2007). Alkaline extraction of membrane fractions was performed to separate membrane and soluble forms of proteins following centrifugation to determine if TMTC3 and TMTC4 are integral membrane proteins (Mostov *et al.*, 1981).

HEK293T cells were transfected with either TMTC3 or TMTC4 S-tag, and proteins were radiolabeled with [^35^S]-Met/Cys for 1 hr. Cells were homogenized in isotonic buffer and fractions were separated by centrifugations. TMTC3 and TMTC4 were found in the nuclear (N) and total membrane (TM) fractions (Fig. 2A, lanes 2 and 4). The nuclear localization of TMTC3 and TMTC4 is likely explained by the contiguous nature of the ER and nuclear membrane prohibiting their separation as the ER proteins calnexin and calreticulin were also in the nuclear fractions. Alkaline extraction of the total membrane fractions followed by centrifugation found TMTC3 and TMTC4 exclusively in the membrane pellet (P) (Fig. 2A, lane 6). This profile was observed for the ER membrane protein, calnexin, and not its soluble paralogue calreticulin, which largely accumulated in the supernatant (S) (Fig. 2A, lane 5). Therefore, both TMTC3 and TMTC4 are integral membrane proteins.

**Figure 2.**
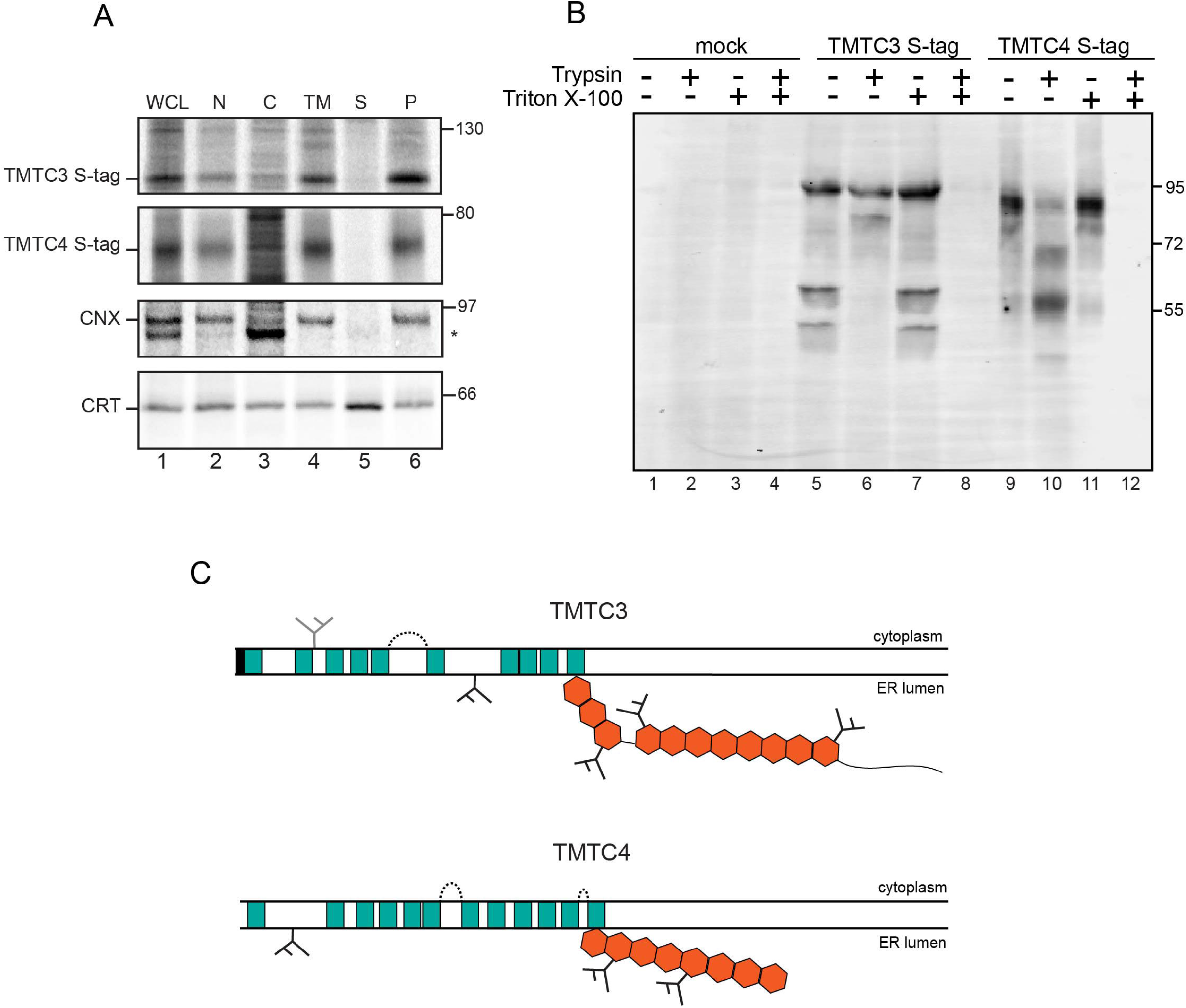
TMTC3 and TMTC4 are transmembrane proteins with their TPR domains facing the ER lumen. (A) HEK293T cells were transfected with S-tagged TMTC3 or TMTC4 as indicated and radiolabeled for 1 hr with [^35^S]-Cys/Met. Cells were homogenized and fractionated prior to alkaline extraction. The fractions collected were whole cell lysate (WGL), nucleus (N), cytosol (C), total membrane (TM), as well as supernatant (S) and pellet (P) fractions upon alkaline extraction of the TM. All samples were subjected to affinity purification with S-tag agarose or calnexin (CNX) and calreticulin (CRT) immunoprecipitation where indicated. Samples were analyzed via SOS-PAGE and detected via autoradiography. (B) TMTC3 S-tag and TMTC4 S-tag were expressed in HEK293T cells. Cells were homogenized and microsomes were isolated by ultracentrifugation then resuspended in homogenization buffer. Aliquots of the ER microsomes were incubated for 15 min at 27 °C with or without triton X-100 and trypsin where indicated. Samples were separated on 9% SOS-PAGE and immunoblotted for the S-tag epitope. (C) Putative trypsin protease cleavage areas (dashed line) on TMTC3 and TMTC4 based on exposed potential cleavage residues and size of cleaved bands in B. Orange hexagons represent ER lumen facing TPR motifs.

Since TMTC3 and TMTC4 are membrane proteins, a trypsin protection assay was employed to determine if their C-terminal TPR motifs are positioned in the ER lumen or the cytoplasm. HEK293T cells were transfected with either TMTC3 or TMTC4 constructs containing C-terminal S-tags, and cells were homogenized prior to isolation of ER-enriched microsomes. Isolated microsomes were resuspended in an isotonic buffer, and aliquots were treated with triton X-100 and trypsin as indicated. Trypsin treatment produced a discrete TMTC3 fragment of ∼83.5 kDa and TMTC4 fragments of ∼67.8 and 57.1 kDa (Fig. 2B, compare lanes 5 to 6 and 9 to 10). As the TPR domains and S-tags are both at the C-termini, this demonstrated that the TPR domains of TMTC3 and TMTC4 were positioned in the ER lumen (Fig. 2C). Combined with the modification of the glycosylation sites added to the TPR rich regions (Fig. 1B), these results demonstrated that the TPR motifs for both of the membrane proteins, TMTC3 and TMTC4, were facing the ER lumen.

Characterization of *TMTC3* and *TMTC4* transcripts was performed to compare TMTC3 and 4 to known O-mannosyltransferases. Both the basal mRNA levels of the two classes of proteins were examined, as well as whether the *TMTC3* and *TMTC4* genes are transcriptionally regulated by ER stress as proteins that reside in the secretory pathway are frequently transcriptionally upregulated by stress (Walter and Ron, 2011). *TMTC1-4* and *POMT1/2* mRNAs were similarly expressed with *TMTC4* and *POMT1* showing the highest levels at ∼2.5% of the reference gene β-actin (Supplemental Fig. 2A). All *TMTC*s and *POMT*s were significantly less expressed than the ER resident chaperone, *BiP*, which is ∼20% of β-actin (Supplemental Fig. 2A).

To determine whether the *TMTC3* and *TMTC4* genes are transcriptionally regulated by ER stress, HEK293A cells were exposed to different ER stress conditions. Cells were subjected to N-glycan synthesis inhibition (tunicamycin), calcium depletion (thapsigargin), redox stress (dithiotreitol, DTT), inhibition of anterograde protein trafficking (brefeldin A) or proteasomal inhibition (MG132). RNA was harvested from cells followed by reverse transcription to generate cDNA and changes in gene expression were measured by qRT-PCR. *TMTC3* gene expression was increased with thapsigargin and brefeldin A treatment by ∼2.1 and 2.5-fold, respectively (Supplemental Fig. 2B). *TMTC4* gene expression was increased with tunicamycin treatment by 1.8-fold and 1.5-fold with brefeldin A treatment (Supplemental Fig. 2B). Thapsigargin, DTT and MG132 did not produce a significant increase in gene expression of *TMTC3*, although the transcription of *BiP* was stimulated by all treatments. Out of the TMTCs, *TMTC4* was up-regulated by stress most significantly and *POMT1* exhibited similar up-regulation in response to stress, which may correspond to their basal mRNA levels being higher or UPR induction upon their respective deletion (Jonikas *et al.*, 2009; Li *et al.*, 2018).

### TMTC3 rescues O-mannosylation of E-cadherin

TMTC1, TMTC2 and TMTC4 have been shown to associate with SERCA2B and play a role in calcium regulation (Sunryd *et al.*, 2014; Li *et al.*, 2018). To investigate the cellular role of TMTC3, initially its binding to SERCA2B was tested. HEK293T cells were transfected with either *TMTC1, TMTC2, TMTC3* or *TMTC4* cDNA. Upon cell homogenization and total membrane isolation, the TMTCs were affinity purified, and associated proteins were resolved by SDS-PAGE. Binding to SERCA2B was monitored by immunoblotting (Fig. 3A). As previously shown, TMTC1, TMTC2 and TMTC4 associated with SERCA2B; however, no interaction between TMTC3 and SERCA2B was observed (Fig. 3A, lane 9). Therefore, unlike the other TMTCs, TMTC3 does not appear to associate with SERCA2B.

**Figure 3.**
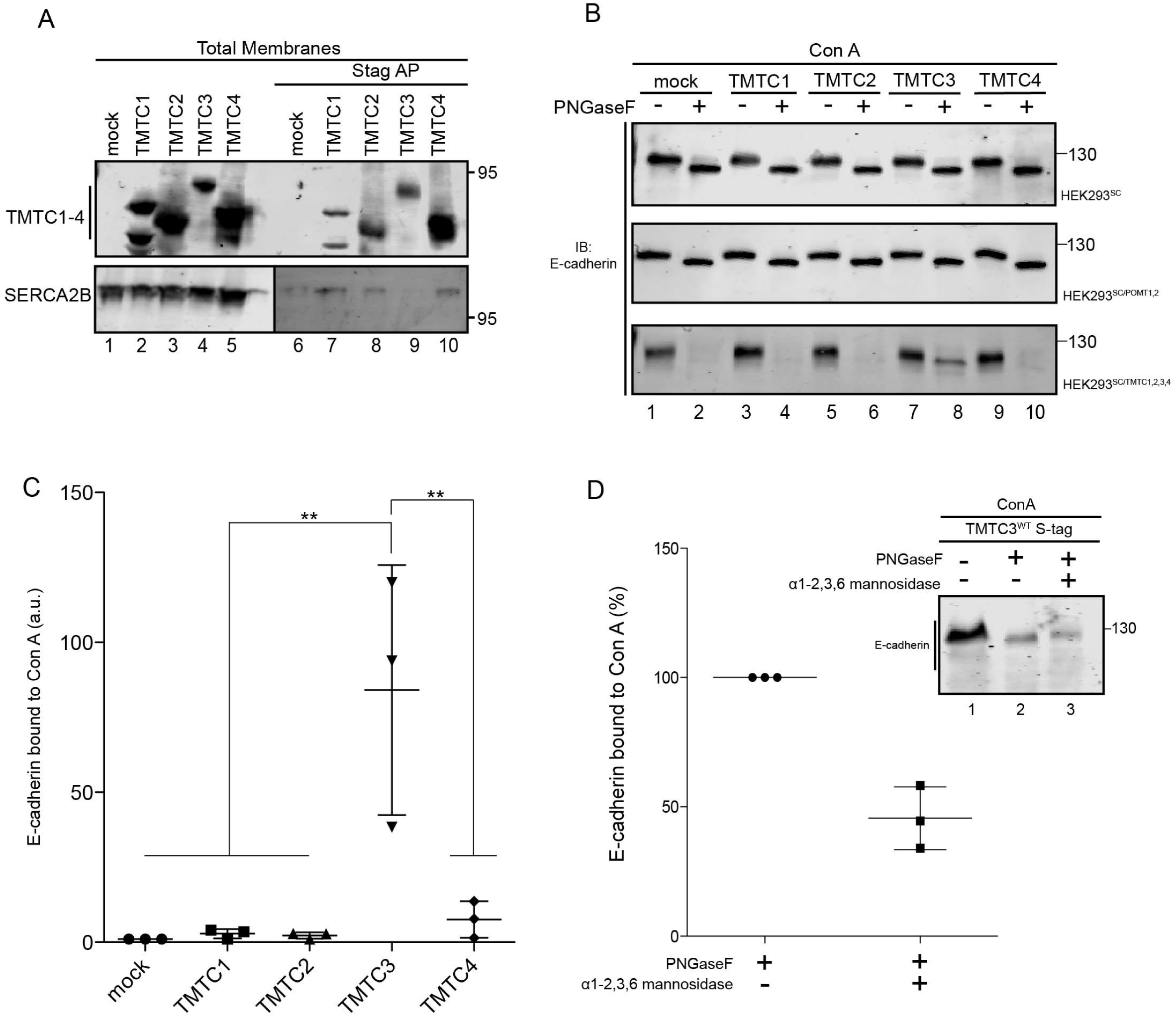
TMTC3 rescues O-mannosylation of E-cadherin in HEK293^SC/TMTC1,2,3,4^ cells. (A) HEK293T cells expressing TMTC1, 2, 3 or 4 were harvested in isotonic buffer and homogenized. A portion of the cell homogenate was subjected to ultracentrifugation and resuspended in reducing sample buffer. This was considered the total membrane fraction (lanes 1-5). An excess of MNT lysis buffer was added to an equal amount of cell homogenate and subjected to S-protein agaraose affinity purification (lanes 6-10). Proteins were detected by immunoblotting with appropriate antisera directed against the S-tag epitope and SERCA2B. (B) S-tagged TMTC1, 2, 3 or 4 cDNA was transfected into indicated cell lines. Cell lysates were collected and split, one half subjected to treatment with endoglycosidase PNGaseF (lanes 2, 4, 6, 8 and 10), prior to pulling down glycosylated proteins with concanavalin A (Con A). Samples were then analyzed by 6% SDS-PAGE and immunoblotted with E-cadherin antisera to assess levels of glycosylated E-cadherin. (C) Quantification of relative density of O-glycosylated E-cadherin from *HEK293*^*SC/TMTC1,2,3,4*^ cells transfected with TMTC1, 2, 3, and 4 cDNA, respectively. Statistical significance between non-transfected cells and TMTC1, 2, 3 or 4 transfected cells was calculated by using one-way ANOVA. Measurements designated (**) have a P value of <0.01. Error bars represent standard deviation. (D) *HEK293*^*SC/TTMTC1,2,3,4*^ cells were transfected with S-tagged TMTC3 cDNA. Cell lysates were collected and subjected to a glycosylation assay with either PNGaseF (lane 2) or combined PNGaseF and a1-2,3,6 mannosidase treatment (lane 3) prior to affinity purification of glycosylated proteins by Con A. Samples were then analyzed via 9% SDS-PAGE and immunoblotted with E-cadherin antisera. Error bars represent standard deviation.

*TMTC1-4* were discovered to contribute to the O-mannosylation proteome when a subset of substrates remained O-mannosylated in cells that lacked the known O-mannosyltransferases, *POMT1* and *POMT2* (Larsen *et al.*, 2017a, 2017b). A significant difference in O-mannosylation of the cadherin family of proteins was observed upon knockout of all four *TMTC*s in HEK293 cells (HEK293^SC/TMTC1,2,3,4^) (Larsen *et al.*, 2017a). Glycoproteomics was used to determine that E-cadherin was O-mannosylated at nine sites (Supplemental Fig. 1A). These results were obtained using HEK293 cells also lacking *COSMC* and *POMGnT1* termed “SimpleCell” or HEK293^SC^ cells. COSMC is an essential chaperone to T-synthase that modifies glycoproteins with an O-GalNAc linked saccharide, and POMGnT1 is a glycosyltransferase that appends β1,2-GlcNAc to O-mannose in the Golgi (Yoshida *et al.*, 2001; Wang *et al.*, 2010). The use of HEK293^SC^ cells created truncated, more homogenous O-mannosylated modifications amenable for glycomics analysis (Vester-Christensen *et al.*, 2013).

To assess potential O-mannosyltransferase activity of the individual TMTCs, a glycosylation and carbohydrate binding assay was employed in glycosyltransferase deficient cell lines including HEK293^SC/TMTC1,2,3,4^ cells with TMTCs complementation (Fig. 3B). These cells were transfected at similar levels individually with S-tagged TMTC1, 2, 3 or 4 (Supplemental Fig. 3A). Cell lysates were collected and divided into two fractions. One fraction of the lysate was treated with PNGaseF to remove N-linked glycans (Fig. 3B, lane 2) prior to both fractions being affinity purified using the lectin, concanavalin A (Con A). Samples were analyzed via SDS-PAGE and immunoblotted with E-cadherin antisera.

E-cadherin levels were not affected by the deletion of COSMC or the glycosyltransferases (Supplemental Fig. 4A). E-cadherin appears to receive O-linked glycans as it was pulled down by Con A even though the glycoproteins lacked N-linked glycans in the PNGaseF treated sample (Fig. 3B, top blot, lane 2). Subsequently, the same assay was performed in HEK293^SC/POMT1,2^ cells and O-glycosylated E-cadherin still bound to Con A (Fig. 4B, lane 2, middle blot). However, in HEK293^SC/TMTC1,2,3,4^ cells, O-glycosylated E-cadherin was not recovered by Con A after N-linked glycan removal suggesting a significant loss of both N- and O-linked glycans (Fig. 3B, lane 2, bottom blot). This result was similarly observed in HEK293^SC/TMTC1,2,3,4^ cells transfected with S-tagged TMTC1, 2 or 4 (Fig. 3B, lanes 4, 6 and 10, bottom blot). However, cells transfected with TMTC3 were able to recover Con A-bound O-linked glycosylated E-cadherin (Fig. 3B, lane 8, bottom blot). The recovery of O-glycosylated E-cadherin by TMTC3 expression was repeated and quantified to confirm that TMTC3 contributed to the O-linked glycosylation of E-cadherin (Fig. 3C).

**Figure 4.**
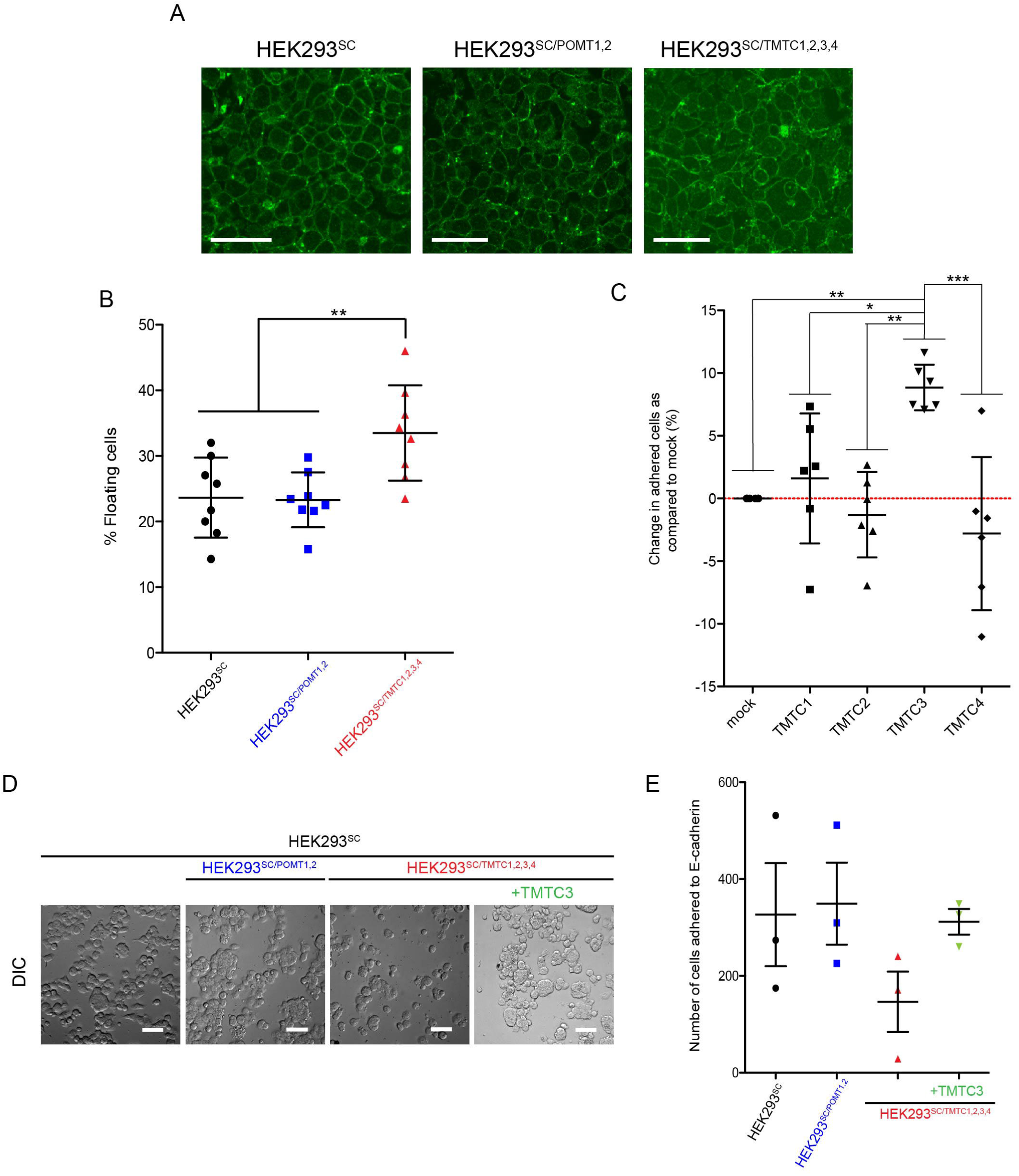
TMTC3 enhances E-cadherin mediated cellular adherence. (A) Cellular localization of cadherins was investigated by confocal microscopy. HEK293^SC^, HEK293^sc/POMT12^ and HEK293^SC/TMTC1,2,3,4^ cells were fixed and stained with pan-cadherin antisera. Scale bars correspond to 50 pm. (B) Adherence was assessed in previously described cell lines. Cells were resuspended in growth media and 1 × 10^6^ cells were subsequently plated in a 24-well plate. Cells were allowed to adhere for 30 min. Unadhered cells were collected and adhered cells were trypsinized and collected. The data describes the % of floating (unadhered) cells over the total (unadhered + adhered) after counting both fractions. Statistical significance between cell types was calculated by using one way ANOVA. Measurements designated (**) have a P value of <0.01. Error bars represent the standard deviation. (C) TMTC1, 2, 3 or 4 cDNAwas transfected into HEK293^SC/TMTC1,2,3,4^ cells and adherence was assessed as described in B. The increase or decrease of adherence of cells transfected with TMTC1, 2, 3 or 4 was established by subtracting the % of mock transfected adhered cells (red dashed line) from the % of adhered, transfected cells (adhered/ (unadhered + adhered)). Error bars represent the standard deviation. Statistical significance between cell treatments was calculated by using one-way ANOVA. Measurements designated *, ** or *** have a P value of <0.05, <0.01 or <0.001, respectively. (D) Cellular adherence to immobilized E-cadherin was assessed via immunoassay. HEK293^SC^, HEK293^sc/POMT1,2^ and HEK293^SC/TMTC1,2,3,4^ cells and HEK293^SC/TMTC1,2,3,4^ cells transfected with TMTC3 were resuspended and 60,000 cells were placed on immobilized E-cadherin (50 μg/mL). Cells were allowed to attach for 1 hr prior to fixing with PFA, permeabilization with triton X-100 and staining with Hoescht. Hoescht nuclear staining and DIC images (pictured) were acquired at 40X magnification. Scale bars correspond to 50 pm. (E) The number of cells adhered to E-cadherin in previously described cell lines was estimated by averaging 5 randomized images capturing nuclei intensity at 405 nm within each well from three independent experiments. Error bars represent the standard deviation.

To determine whether TMTC3 mediated O-linked glycosylation of E-cadherin included O-mannosylation, a combinatorial endoglycosidase assay was performed. S-tagged TMTC3 was transfected into HEK293^SC/TMTC1,2,3,4^ cells and the lysate was subsequently treated with PNGaseF or both PNGaseF and α1-2,3,6 mannosidase prior to affinity purification with Con A, as the mannosidase would specifically remove mannose residues. O-glycosylated E-cadherin was recovered from cells complemented with TMTC3 after PNGaseF treatment; however, the amount bound to Con A upon subsequent α1-2,3,6 mannosidase treatment was reduced by ∼45%, indicating that TMTC3 contributes to O-mannosylation of E-cadherin (Fig. 3D insert, compare lanes 3 to 2).

### TMTC3 enhances cellular adherence and binding to E-cadherin

Studies on protein O-mannosylation in yeast have revealed that O-glycans are important for protein stability, trafficking, localization, function and stress response (Yang *et al.*, 2009; Petkova *et al.*, 2012; Xu and Ng, 2015). To understand the effect of TMTC3 on E-cadherin, the expression, stability and trafficking of E-cadherin was assessed in *TMTC1-4* knockout cells. Briefly, HEK293^SC^, HEK293^SC/POMT1,2^ and HEK293^SC/TMTC1,2,3,4^ cells were lysed and subjected to lectin affinity purification with Con A prior to samples being analyzed via SDS-PAGE and immunoblotted with E-cadherin antisera. There was no significant difference observed in expression between cell types (Supplemental Fig. 4A). Cellular trafficking was also assessed by endoglycosidase sensitivity in combination with cycloheximide chase and a significant difference was not observed (data not shown). Cell surface localization of several cadherins, including E-cadherin, was monitored using confocal immunofluorescence microscopy. HEK293^SC^, HEK293^SC/POMT1,2^ and HEK293^SC/TMTC1,2,3,4^ cells were fixed and stained for pan-cadherin (Fig. 4A). In all three cell lines, cadherins localized to the cell surface. Interestingly, a morphology defect was observed in cells lacking *TMTC1-4*. While cell surface staining in HEK293^SC^ and HEK293^SC/POMT1,2^ cells appeared round and evenly distributed, cadherin staining in HEK293^SC/TMTC1,2,3,4^ cells showed irregularity.

Cell-cell adhesion interactions are essential for embryogenesis, tissue morphogenesis and renewal (Halbleib and Nelson, 2006). Adherins junctions are the sites of cell-cell contact where adhesion is mediated by cell-surface receptors of the cadherin family (Meng and Takeichi, 2009). E-cadherin plays a critical role in cell-cell adhesion and development and O-mannosylation has been shown to be important for its role in forming adherins junctions (Lommel *et al.*, 2013). To determine the effect of TMTCs on E-cadherin mediated cellular adherence, general cellular adherence was evaluated. HEK293^SC^, HEK293^SC/POMT1,2^ and HEK293^SC/TMTC1,2,3,4^ cells were re-suspended in growth media and subsequently plated and allowed to adhere for 30 min. After 30 min, the floating cells were collected for each cell type and adhered cells were trypsinized and collected. Each fraction was quantified and the percent of floating cells over the total was calculated. HEK293^SC^ and HEK293^SC/POMT1,2^ cells showed similar levels of adherence with average floating cell percentages at ∼23% (Fig. 4B). Contrastingly, HEK293^SC/TMTC1,2,3,4^ cells demonstrated reduced cellular adherence as the percentage of floating cells observed was ∼33% (Fig. 4B). To determine if any one of the TMTCs individually contributed to cellular adherence *TMTC1, 2, 3* and *4* cDNA was transfected into HEK293^SC/TMTC1,2,3,4^ cells. The adherence assay described above was performed and the percentage of adhered cells as compared to mock was calculated. Upon establishing a baseline using the mock treated cells, TMTC3 transfected cells were able to recover adherence the most with an observed increase over mock of ∼9% (Fig. 4C).

Cadherins are cell-surface membrane glycoproteins that contain multiple repeats of extracellular cadherin domains and they mediate cell-cell adhesion by trans homodimerization between apposed cells (Cavallaro and Christofori, 2004; Yagi, 2008; Brasch *et al.*, 2012). E-cadherin mediated cell-cell adhesion was assessed in HEK293^SC^, HEK293^SC/POMT1,2^ and HEK293^SC/TMTC1,2,3,4^ cells to determine if genetic manipulation of the glycosyltransferases affected its adhesion function. Purified extracellular domain of E-cadherin was incubated overnight to allow for attachment to the plate surface. HEK293^SC^, HEK293^SC/POMT1,2^ and HEK293^SC/TMTC1,2,3,4^ cells were then resuspended and ∼60,000 cells were seeded on the purified substrate. Cells were allowed to adhere for 1 hr prior to discarding unadhered cells. Adhered cells were then fixed with paraformaldehyde before permeabilization with triton X-100, and nuclei were stained with Hoescht. Hoescht images were captured at five different locations within each well and the number of nuclei quantified. HEK293^SC/TMTC1,2,3,4^ cells showed markedly reduced binding to the extracellular region of E-cadherin compared to HEK293^SC^ and HEK293^SC/POMT1,2^ cells (Fig. 4D and E). The same assay was performed on HEK293^SC/TMTC1,2,3,4^ cells expressing TMTC3 and E-cadherin mediated adherence was recovered. These results, combined with the results from the carbohydrate binding assay (Fig. 3B and C), suggest that TMTC3 affects E-cadherin mediated adherence through O-mannosylation of the glycoprotein.

### TMTC3 disease variants exhibit reduced stability

Cell-cell adhesion interactions are essential for embryogenesis, tissue morphogenesis and renewal, all key processes of development (Halbleib and Nelson, 2006). Cobblestone lissencephaly is a severe brain malformation characterized by irregular borders, dysplasia, hypoplasia and dysmyelination, which are due to overmigration of neurons and glial cells beyond the external basement membrane. The cause of this over migration defect is impaired interactions between glial limitans and the extracellular matrix of the basement membrane (Siegenthaler and Pleasure, 2011; Barkovich *et al.*, 2012). Glycosylated cell surface binding receptors provide a physical link between the cytoskeleton of the glial cells and the basement membrane (Barresi and Campbell, 2006). Loss of glycosylation of these molecules contributes to functional defects during development by reducing binding to the extracellular matrix (Michele *et al.*, 2002). Cobblestone lissencephaly has been associated with mutations in sixteen genes, and twelve of these genes are involved in O-linked glycosylation, including *POMT1* and *2* (Jerber *et al.*, 2016).

Recently, a study identified biallelic mutations in *TMTC3* from Cobblestone lissencephaly patients exhibiting intellectual disability, hypotonia and delayed milestones (Jerber *et al.*, 2016). Six families had a total of eight distinct mutations in affected individuals, four of which were homozygous and two were compound heterozygous. Using predictive algorithms, the altered organization of seven of these mutants and a compound heterozygous mutant (R71H) associated with another neuronal disorder (PVNH) was analyzed (Supplemental Fig. 5A). The point mutations do not appear to alter organization; however, the resulting residue mutations change the charge of the native residue, which may have some topological or functional effect on TMTC3 as some of these mutations are in close proximity to the putative active sites (Supplemental Fig. 5A, H67D, R71H and G384E). The frame shift mutations beginning at residue positions 488, 562 and 654 greatly change the number and organization of the predicted TPR motifs resulting in truncated forms of TMTC3. Additionally, these frameshift mutations could result in five to seven non-native amino acids being transcribed before the premature stop codon. The frameshift mutation at position 841 results in a shortened C-terminus and three putative non-native residues, and the early stop codon mutation at position 873 would result in a protein that is missing 41 amino acids.

TMTC3 disease variants were characterized for ER localization via glycosidase assay and immunofluorescence confocal microscopy. TMTC3 disease variant constructs, engineered to harbor any resulting non-native residues and C-terminal S-tags, were transfected into HEK293T cells prior to cell lysate and media fractions being affinity purified with S-protein agarose beads followed by glycosidase treatment. Shifts upon PNGaseF treatment (Supplemental Fig. 5B, lanes 3 and 9) were observed for all constructs indicating that these proteins were targeted to the ER and received N-linked glycans. A similar increase in mobility was observed upon Endo H treatment indicating that the carbohydrates were high mannose glycoforms (Supplemental Fig. 5B, lanes 2 and 8). All TMTC3 disease variants co-localized with calreticulin (CRT), while co-localization was not observed with the Golgi marker, GM130, when immunofluorescence staining was performed in transfected COS7 cells (Supplemental Fig. 6). Therefore, the glycosylation profiles and the cellular distribution are consistent with TMTC3 disease variants residing in the ER like wild type (WT) TMTC3.

The only exception to ER localization studies was the frameshift mutation R488Efs. This mutant did not express in COS7 cells so cellular localization via immunofluorescence was unable to be determined. In order to assess expression of R488Efs S-tag, MG132, a proteasome inhibitor, was added to the media of cultured cells transfected with the S-tagged mutant construct. The immunoblot revealed that R488Efs was stabilized by proteasome inhibition suggesting that once translated, it is quickly degraded (Fig. 5C, compare lane 3 to 4). Additionally, a glycosylation profile was obtained using proteasome inhibition (Supplemental Fig. 5B, top right panel). A shift upon EndoH and PNGaseF treatment was not observed, therefore this mutant may not be glycosylated. This could be explained by the predictive organizational outcome of this variant as the frameshift results in a severe truncation of the protein, including most of the predicted N-linked glycosylation sites (Supplemental Fig. 5A).

**Figure 5.**
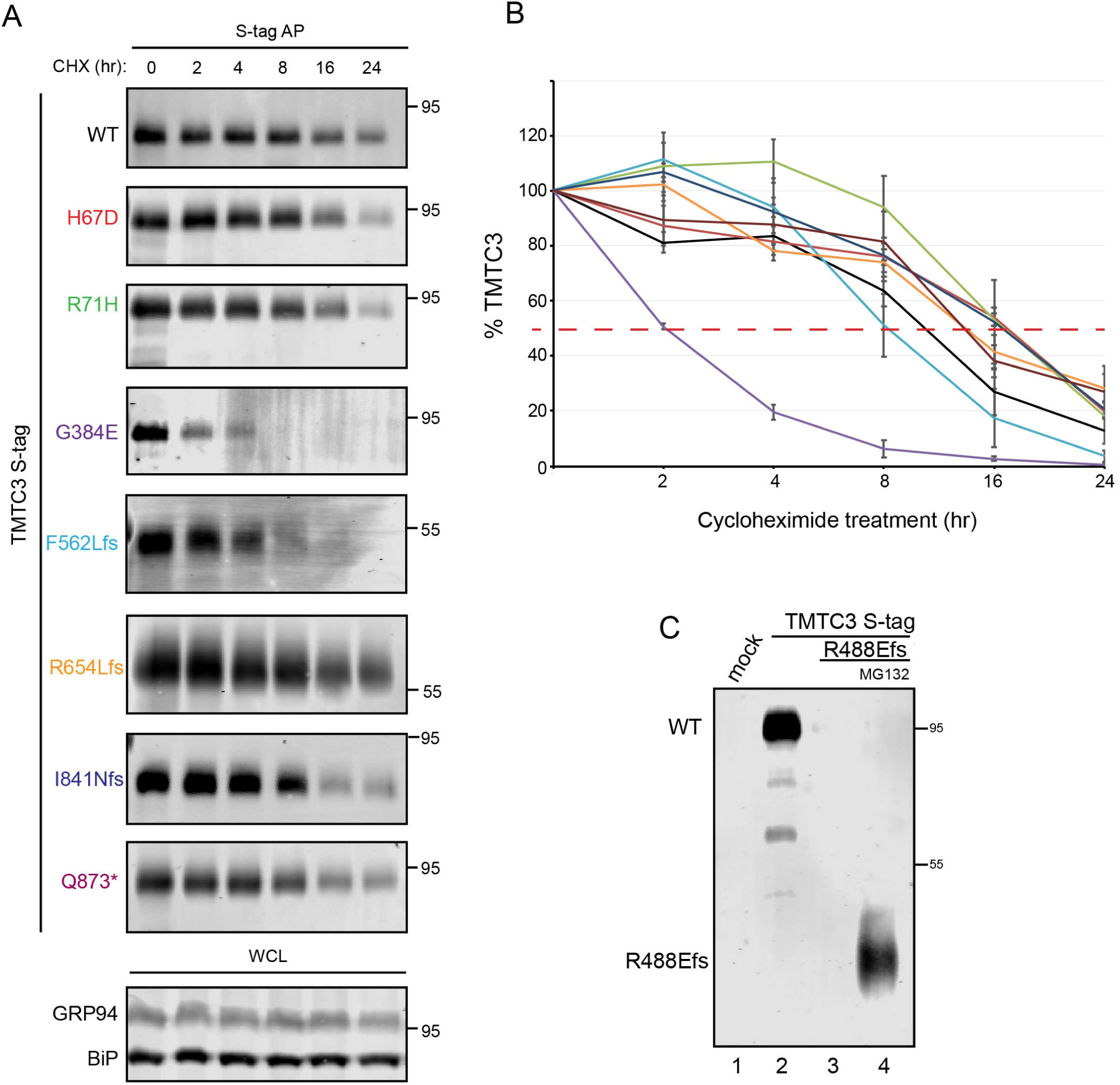
TMTC3 WT and disease variant stability. (A) HEK293T cells were transfected with S-tagged TMTC3 WT or disease variant cDNA as indicated. Cells were treated with 100 μg/ml cycloheximide for the indicated time prior to collection, lysed in MNT and samples were subjected to affinity purification with S-protein agarose. Reducing sample buffer was added and samples were analyzed via 9% SDS-PAGE and immunoblotted for the S-tag epitope. Cell lysates were also subjected to precipitation with 10% trichloroacetic acid, analyzed via 9% SDS-PAGE and immunoblotted with KDEL antisera to determine stability of GRP94 and BiP (WCL). (B) Time points were collected at 0, 2, 4, 8, 16 and 24 hr after cycloheximide treatment. The amount of TMTC3 protein remaining at 2, 4, 8, 16 and 24 hr was quantified and normalized to the starting material (0 h) and averaged from three independent experiments. Error bars represent standard error of the mean. (C) HEK293T cells were transfected with S-tagged TMTC3 WT or the TMTC3 R488Efs disease variant as indicated. Cells transfected with the TMTC3 R488Efs variant were treated with 20 μM MG132 for 12 hr prior to collection and lysis in MNT (lane 4). Samples were subjected to affinity purification with S-protein agarose and analyzed via SOS-PAGE prior to immunoblotting for the S-tag epitope.

The rapid degradation of the R488Efs variant suggested that instability of TMTC3 could lead to disease, therefore the stability of the remaining disease variants was analyzed. HEK293T cells were transfected with the S-tagged TMTC3 mutant constructs and cells were treated with cycloheximide for 0-24 hr. Cells were collected at the indicated time points, lysed and subjected to S-protein affinity purification (Fig. 5A). WT TMTC3 has a half-life of ∼12 hr (Fig. 5B, black line). Several of the mutants show similar or slightly extended half-lives; however, F562Lfs and G384E mutations lead to significantly reduced half-lives of ∼8 and 2 hr, respectively (Fig. 5B, light blue and purple lines). This increased instability could explain a disease phenotype. Three of the disease variants are rapidly recognized as non-native and degraded. Further biochemical investigation is necessary to better understand the effects of such mutations.

### TMTC3 and TMTC4 knockdown delays gastrulation in Xenopus laevis

Tissue morphogenesis during development is dependent upon the cadherin family of cell-cell adhesion molecules, which includes classical cadherins, protocadherins and atypical cadherins (Halbleib and Nelson, 2006). During vertebrate morphogenesis, the various cadherin family members are differentially expressed in a variety of tissues (Nandadasa *et al.*, 2009). E-cadherin has been shown to play a central role in cell-cell adhesion during embryo development before implantation. If O-mannosylation is prevented either genetically or biochemically, embryo development is arrested (Lommel *et al.*, 2013). Staining for E-cadherin during mouse embryonic development showed that it is retained basolaterally, however, it was reduced at sites that also showed reduced adherence.

Analysis of the TMTCs protein sequence conservation among commonly studied species revealed strong similarity to the Tmtc paralogues in *Xenopus laevis*, therefore the potential phenotypic effects of Tmtc knockdown in developing *Xenopus* embryos was assessed (Supplemental Fig. 7A). In *Xenopus*, C-cadherin is the major cadherin expressed maternally persisting during early cell division and gastrulation. During gastrulation, E-cadherin is expressed in the ectoderm. At the end of gastrulation, the ectoderm becomes segregated into the dorsal neural ectoderm, which then activates expression of N-cadherin and turns off E-cadherin. The non-neural ectoderm retains E-cadherin expression becoming the epidermis (Nandadasa *et al.*, 2009). A global mRNA expression study conducted in developing *Xenopus* embryos showed that *tmtc3* and *tmtc4* mRNA are maternally expressed and that *tmtc3* is strongly expressed during the mid-blastula transition (MBT) and remains abundant throughout gastrulation (Session *et al.*, 2016)(Supplemental Fig. 7B). As both *e-cadherin* and *tmtc3* and *tmtc4* mRNA begin to increase in expression prior to gastrulation phase, the effect of Tmtc knockdown was observed during gastrulation.

Morpholino sequences (Genetools LLC) were designed to match the ATG region of *Xenopus laevis* Tmtc1 to 4 to block protein translation of both allo allele (L and S). Individual morpholinos were microinjected into embryos at the 1-cell stage to ensure that every cell contained morpholino upon subsequent divisions. The injected embryos were allowed to develop as normal and gastrulation was observed via time-lapse video microscopy using an inverted light microscope. The diameter of the blastopore was measured during gastrulation (stage 10-12) for all injected embryos as well as non-injected controls. Those injected with the Tmtc1, 2, 3 or 4 morpholino showed a significant delay in gastrulation as exhibited by a larger blastopore diameter than non-injected controls at the same time point (Fig. 6A and B). The most significant effect was observed for Tmtc3 and Tmtc4 morpholino injected embryos. With *tmtc3* mRNA showing the highest expression in development (Supplemental Fig. 7B) and to confirm specificity of this delay in gastrulation, human *WT TMTC3* mRNA was subsequently injected after the Tmtc3 morpholino in 1-cell stage embryos and gastrulation was observed as previously described (Fig. 6A and B). Human WT TMTC3 was able to completely rescue the delay in gastrulation caused by knocking down *Xenopus* Tmtc3, suggesting that the developmental defect is specific to TMTC3 and that the human and *Xenopus* proteins share similar functions.

**Figure 6.**
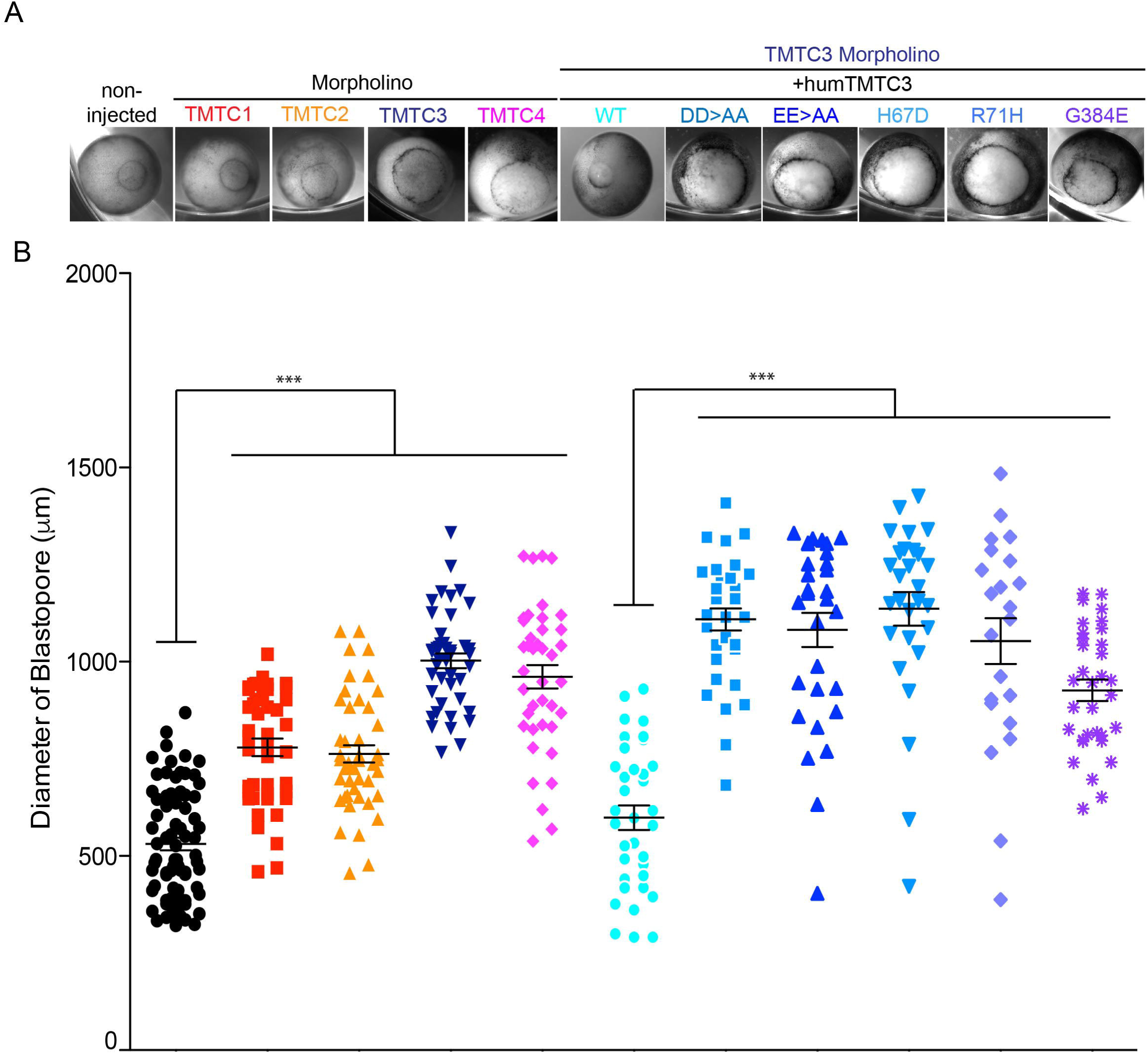
Knockdown of Tmtc3 in developing Xenopus laevis emrbyos affects gastrulation. (A) Embry-os were injected with 10 ng of morpholinos against Tmtc1, Tmtc2, Tmtc3 and Tmtc4 at the one-cell stage. The embryos were scored for gastrulation when control embryos reached stage 11-12. Embryos were also injected with 10 ng of Tmtc3 morpholino and subsequently injected with S-tagged huTMTc3^WT^, huTMTC3^DD>AA^ huTMTC3^EE>AA^, huTMTC3^H67D^, huTMTC3^R71H^ and huTMTC3^G384E^ at the one-cell stage. The embryos were scored as described above. (B) The average of three or more independent experiments is plotted on the graph. Statistical significance between injection conditions was calculated using one-way ANOVA. Measurements designated(***) have a P value of <0.001. The error bars represent the standard deviation to the mean. The number of embryos analyzed are as follows: non-injected n=75, TMTC1 MO n=43, TMTC2MO n=45, TMTC3MO n=43, TMTC4MO n=40, TMTC3MO + huTMTC3 n=35, TMTC3MO + huTMTC3^DD>AA^ n=33, TMTC3MO + huTMTC3^EE>AA^ n=30, TMTC3MO + huTMTC3 ^H67D^ =28, TMTC3 MO + huTMTC3 ^R71H^ n=22 and huTMTC3^G384E^ n =33.

The O-mannosyltransferases POMT1/2 and Pmt1-7 contain diacidic motifs positioned near the membrane in the lumen of the ER, which are hypothesized to serve as the active site for the transferase activity (Supplemental Fig. 1B) (Lommel *et al.*, 2011; Bai *et al.*, 2019). TMTC3 has two N-terminal diacidic motifs that are proposed to be luminally orientated, Asp31Asp32 (DD) and Glu63Glu64 (EE). (Supplemental Fig. 1C). To determine whether a putative active site could be assigned to TMTC3, the diacidic motifs were mutated to di-Ala motifs. The TMTC3 DD>AA and EE>AA constructs were injected at the 1-cell stage with Tmtc3 morpholino. While injected embryos were observed for gastrulation, a number of embryos were also collected to assess construct expression. Briefly, embryos were homogenized in lysis buffer prior to total protein precipitation with trichloroacetic acid and analysis by SDS-PAGE and immunoblotting for S-tag epitope (Supplemental Fig. 7C). The analysis showed that all constructs, human WT TMTC3, DD>AA and EE>AA were similarly expressed. While the WT TMTC3 was able to rescue the gastrulation defect, neither the DD>AA nor EE>AA construct could (Fig. 6A and B). Given that a simple point mutation in TMTC3 (G384E) resulted in significantly decreased stability of the protein (Fig. 5A and B), it is possible that mutating the two negatively charged residues within the first luminal loop of TMTC3 to neutral Ala could affect function of TMTC3.

Mutations in *TMTC3* have been identified in patients with diseases resulting from defects in cell migration. Since the process of gastrulation in metazoans is dependent upon a series of complex and coordinated cell movements, TMTC3 disease variants were assessed in the previously described *Xenopus* gastrulation assay (DeSimone *et al.*, 2007; Winklbauer, 2012). One-cell stage embryos injected with Tmtc3 morpholino were subsequently injected with human *TMTC3* mRNA harboring three of the point mutations found in patients, H67D, R71H and G384E (Supplemental Fig. 5A). The embryos were allowed to develop to gastrulation prior to being assessed as previously described. Tmtc3 morpholino injected embryos display an average increased blastopore diameter of ∼880 μm compared to non-injected or WT TMTC3 rescue controls, ∼530 and ∼600 μm, respectively (Fig. 6). TMTC3 disease mutants H67D, R71H and G384E possessed increased blastopore diameters of ∼1100, ∼1000 and ∼930 μm, respectively. This data shows that while WT TMTC3 rescues the gastrulation defect, the disease variants resulting in point mutations in TMTC3 increased developmental delays suggesting that they may act as dominant negative mutants.

## Discussion

Our study demonstrates that TMTC3 contributes to the O-mannosylation of E-cadherin and this modification is important for E-cadherin mediated cell-cell adhesion. TMTC3 is an integral ER membrane protein, likely adopting a polytopic structure, with its putative TPR region residing within the ER lumen. The knockdown of Tmtc3 in *Xenopus laevis* led to a delay in embryonic gastrulation, a process that is heavily dependent on cell-cell adhesion and migration (Winklbauer, 2012). Although the TMTCs all share similar protein domain architecture and the *TMTC* mRNA profiles is comparable to previously known O-mannosyltransferases, POMT1 and 2, TMTC3 appears to play a critical role in cell adhesion. This study provides insight into cell migration diseases caused by mutations in *TMTC3*.

TMTC3 shares 24-28% identity and 38-43% similarity in amino acid sequence with the other members of the TMTC protein family, suggestive of a common evolutionary origin (Sievers *et al.*, 2014). TMTC3 and TMTC4 are present in the chordata phylum and recent analysis shows that they are conserved in protists (Larsen *et al.*, 2019). Like TMTC1 and TMTC2, TMTC3 and TMTC4 are divided into two regions that appear to create distinct domains, an N-terminal hydrophobic region and a C-terminal domain consisting of large TPR clusters (Sunryd *et al.*, 2014). ΔG predicts that TMTC3 and TMTC4 have ten and twelve potential transmembrane domains, respectively (Fig. 1A). Alkaline extraction of total membrane preparations demonstrated that TMTC3 and TMTC4 are integral membrane proteins and trypsin digestion of these preparations produced partially protected C-terminal fragments of both TMTC3 and TMTC4 that correspond to the entire predicted C-terminal TPR region, as well as a number of the hydrophobic segments. Therefore, TMTC3 and TMTC4 appear to be polytopic membrane proteins with their hydrophobic N-termini providing long stretches of multiple hydrophobic segments and exposed cytoplasmic loops.

Protein O-mannosylation involves the transfer of a mannose from an activated dolichol monophosphate mannose to the hydroxyl group on a Ser or Thr. The O-mannosyltransferase activity for yeast Pmts appears to involve a diacidic motif (Asp-Glu) that resides near the membrane in luminal loop 1 of the polytopic proteins (Lommel *et al.*, 2011; Bai *et al.*, 2019)(Supplemental Fig. 1B). TMTC3 possesses two diacidic motifs in the first predicted luminal loop (Asp31Asp32 (DD) and Glu63Glu64 (EE)). These diacidic residues were mutated to Ala and the stability of TMTC3 was monitored (Supplemental Fig. 3B, lower panel). While TMTC3^EE>AA^ was expressed at similar levels to WT TMTC3, TMTC3^DD>AA^ was not visualized after transfection. To determine if TMTC3^DD>AA^ was unstable, a cycloheximide chase was carried out for WT TMTC3, TMTC3^DD>AA^, and TMTC3^EE>AA^ (Supplemental Fig. 3C). TMTC3^DD>AA^ was turned over rapidly when compared to WT TMTC3 and TMTC3^EE>AA^ indicating that TMTC3^DD>AA^ was unstable. Even though the instability of TMTC3^DD>AA^ prohibited us from comparing its ability to rescue the O-mannosylation of E-cadherin in HEK293^SC/TMTC1,2,3,4^ cells to WT TMTC3 and TMTC3^EE>AA^, we were able to show that TMTC3^EE>AA^ rescues O-glycosylation of E-cadherin as E-cadherin still binds Con A in TMTC3^EE>AA^ transfected cells (Supplemental Fig. 3B, lane 8). This potentially links the TMTC3 Asp31Asp32 (DD) diacidic motif to its activity. In *Xenopus*, neither the TMTC3^EE>AA^ nor TMTC3^DD>AA^ construct was able to rescue the delay in gastrulation caused by the Tmtc3 morpholino (Fig. 6A and B). These results indicated that the diacidic residues in loop 1 were important for protein stability, possibly the transferase activity or complex formation.

The glycosylation profile, combined with the trypsin protection of the C-terminal domains, placed the TPRs within the ER lumen for both TMTC3 and TMTC4. This is similar to the TPR regions of TMTC1 and TMTC2 (Sunryd *et al.*, 2014). TMTC3 and TMTC4 encode eleven and eight TPR motifs, respectively, according to TPRPred (Karpenahalli *et al.*, 2007). TPR domains are found in proteins across species and organelles and participate in a number of activities including protein translocation, folding, and post-translational modification (Graham *et al.*, 2019). Furthermore, TPR-containing proteins comprise the largest functional family within the cellular proteostasis/chaperome network (Brehme *et al.*, 2014). TPRPred predicted that the eleven TPR motifs of TMTC3 are organized into clusters of three and eight, while TMTC4 has eight sequential TPRs. The differences in the organization of the TPR domains for TMTC3 and TMTC4 could have implications for their functions.

Extensive bioinformatic analysis shows that long stretches of consecutive TPR domains (observed up to sixteen) tend to be more promiscuous in their selectivity (Magliery and Regan, 2005). Given that TMTC3 and TMTC4 both possess a stretch of eight consecutive motifs, this suggests that they may bind a number of substrates. This is most similar to the well-studied TPR-containing cytoplasmic O-GlcNAc transferase (OGT), which contains a stretch of 12.5 TPRs. The structure of OGT revealed that the TPRs form a super helix with conserved Asn along the concave surface that form a continuous ladder. The Asn in OGT that are particularly well-conserved are those at positions 6 and 9 in the TPR consensus and thought to contribute to inter-repeat interactions (Jínek *et al.*, 2004). Seven of the eleven TPRs in TMTC3, as predicted by TPRPred, have Asn at position six and seven out of eight of the TPRs in TMTC4 have Asn at position six in addition to two of the eight having Asn at position nine. The number of conserved Asn at positions six and nine of the predicted TPRs in TMTC3 and TMTC4 suggests that they may form super helical structures and use the Asn to nucleate protein-protein interactions as this also mimics the Asn conservation observed in the ARM-repeat proteins importin-α and β-catenin, which contribute to the binding of the target peptide (Conti *et al.*, 1998; Huber and Weis, 2001). β-catenin was shown to interact with E-cadherin through conserved Asn on its ARM-repeat domains. β-catenin has twelve ARM-repeat domains and five of the domains contain an Asn at position 13 and are responsible for recognizing the backbone of extended peptides. Furthermore, alignment of murine cadherin family sequences (M-cadherin, OB-cadherin, desmocollin 1a and desmoglein 1) to E-cadherin show that many of the observed E-cadherin/β-catenin interactions are likely to be conserved among other cadherin family members. TMTC3 contains seven Asn at a conserved position on its TPR motifs, which could be used to interact with the well conserved family of cadherins in order to recruit the substrates for O-mannosylation near the ER membrane.

Analysis of TPR-containing protein structures and their ligands revealed that smaller clusters of three and four TPRs corresponds to co-chaperone function (Brehme *et al.*, 2014; Graham *et al.*, 2019). Like one of the most well-studied TPR structures in the protein HoP, the N-terminal cluster of three TPRs in TMTC3 may bind and recruit chaperones for protein homeostasis, analogous to the MIR domains in the POMT/Pmts (Scheufler *et al.*, 2000; Fujimori *et al.*, 2017). MIR domains are observed in six non-redundant proteins, all of which target to the ER (Nielsen, 2017; Letunic and Bork, 2018). Stromal cell-derived factor 2 (SDF2) is a small protein of 211 amino acids comprised primarily of three MIR domains. While not much is known about MIR domain function for SDF2 and its isoform, SDF2L1, their MIR domains have been found to regulate the chaperone cycle of BiP (Fujimori *et al.*, 2017). Both proteins form a stable complex with the ER J-domain protein ERdj3 and inhibit the aggregation of misfolded ER cargo by binding non-native proteins and promoting the BiP-substrate interaction cycle. We hypothesize that the three TPR motif cluster of TMTC3 may form a similar complex with chaperones for the delivery of unfolded substrate in order to expose putative sites for O-mannosylation.

Thirty-seven members of the cadherin superfamily of cell membrane receptors were identified as major carriers of O-mannose glycans (Vester-Christensen *et al.*, 2013). Cell-surface receptors of the cadherin family mediate cell-cell adhesion at sites of cell-cell contact, which is crucial for cells to function in an integrated manner (Lodish *et al.*, 2000; Meng and Takeichi, 2009). In our study, cells lacking *TMTC1-4* stained for cadherins display cell-cell attachment irregularity appearing less round and consistent in shape (Fig. 4A). In combination with the data demonstrating that TMTC3 is involved in the O-mannylation of E-cadherin, this suggests that TMTC3 plays a role in cellular adherence through the O-mannosylation of E-cadherin.

Cadherins contain multiple repeats of extracellular cadherin (EC) domains and they mediate cell-cell adhesion by trans homodimerization between the most distal EC1 and EC1-2 domains on apposed cells (Cavallaro and Christofori, 2004; Yagi, 2008; Brasch *et al.*, 2012). The other EC domains play a critical role in presenting the EC1 and 2 domains so that they may form these homodimers. The O-mannose modified sites identified are confined to the EC2-5 domains of both classic type 1 and 2 cadherins and these sites also appear to be evolutionarily conserved. E-cadherin, a type I classical cadherin, holds epithelial sheets together and is highly abundant at the sites of cell-cell contact along lateral surfaces (Lodish *et al.*, 2000). We found that HEK293^SC/TMTC1,2,3,4^ cells adhere less to the immobilized extracellular region of E-cadherin than HEK293^SC^ and HEK293^SC/POMT1,2^ cells suggesting that genetic manipulation of the *TMTC*s affects E-cadherin’s ability to homodimerize (Fig. 4D and E). Complementation of HEK293^SC/TMTC1,2,3,4^ cells with TMTC3 increased E-cadherin mediated cell-cell adhesion, therefore demonstrating that TMTC3 O-mannosylation of E-cadherin plays a crucial role in its ability to homodimerize (Fig. 4D and E, and Fig. 3). Given that E-cadherin is also crucial during embryogenesis, the effects we observe upon knockdown of Tmtc3 and subsequent rescue of the gastrulation delay with human TMTC3 WT, could be attributed to the role TMTC3 plays in E-cadherin mediated cell-cell adhesion and O-mannosylation (Fig. 6)(Halbleib and Nelson, 2006).

Mutations in *TMTC3* were identified in patients with Cobblestone lissencephaly and PVNH, common brain malformations caused by defects in neuronal migration (Jerber *et al.*, 2016; Farhan *et al.*, 2017). More specifically, three of the mutations (R488Efs, F562Lfs and G384E variants) identified in Cobblestone lissencephaly patients that possess a significant number of clinically observed neurological defects were found to be unstable in our protein degradation assay (Jerber *et al.*, 2016)(Fig. 5). The rapid degradation of these variants could explain the development delays observed in patients due to lack of *TMTC3*. Our *Xenopus* developmental studies revealed that two of the mutants (H67D and R71H) lead to a developmental delay even though they were more stable in cells. Our characterization of the disease variants of TMTC3 provides a link between a previously uncharacterized protein and disease resulting from its mutation.

*TMTC1-4* are not only implicated in the O-mannosylation of cadherins but were also found to be involved in the O-mannosylation of several ER/Golgi resident proteins, including ERp57, ERdj4 and FKBP10 (Larsen *et al.*, 2017a). Each of these are important chaperones for protein folding of the secretory pathway aiding in disulfide bond formation, BiP activation and proline *cis trans* isomerization. O-mannosylation of both trafficked and resident secretory pathway proteins could contribute to their quality control either through increasing favorable folding or enhancing degradation of specific substrates upon misfolding. While previous studies show that O-mannosylation is an important structural modification, recent studies show that it may also play a role in ER quality control (ERQC) (Xu and Ng, 2015). If certain mutants are stabilized, they could cause dysregulation of protein homeostasis due to an excess of O-mannose modification of particular ER resident chaperones.

In conclusion, we demonstrate that TMTC3 is crucial for O-glycosylation of E-cadherin and cell-cell adhesion and embryonic development. Evidence shows that although the TMTCs share architectural similarity, they perform distinct functions and help to regulate cellular homeostasis by participating in essential processes, post-translational modification and calcium regulation. Altogether, TMTC3 is likely an enzyme mediating O-mannosylation, and we have shown that TMTC3 directed O-mannosylation is biologically important, however further studies are needed to better understand TMTC3’s direct activity as an O-mannosyltransferase and whether it requires other proteins for its activity, its function within the cell and how mutations in this gene cause disease.

## Experimental Procedures

### Plasmids and reagents

Dulbecco’s modified Eagle’s medium (DMEM), DMEM GlutaMax, fetal bovine serum, penicillin and streptomycin were purchased from Invitrogen/ThermoFisher Scientific. Easy-Tag [^35^S]-Cys/Met was purchased from PerkinElmer Life Sciences. S-protein-agarose beads and S-tag antibody were purchased from EMD Millipore. Endo H, PNGase F, Protoscript II first strand cDNA synthesis kit and all cloning reagents were purchased from New England Biolabs. FastStart SYBR Green qPCR mix was purchased from Roche Diagnostics, and all primers were acquired from IDT DNA. IRDye® 800CW Goat anti-Mouse IgG and IRDye® 680RD Goat anti-Rabbit IgG were purchased from Li-COR Biosciences. Protein A sepharose CL-4B was purchased from GE Healthcare. Antibodies directed toward the following antigens were also purchased: calnexin (Enzo Life Sciences); calreticulin (ThermoFisher Scientific); E-cadherin (GeneTex); pan-cadherin (Sigma); ERp57 (gift from Dr. Taku Tamura, Akita University, Japan), GM130 (BD Biosciences) and KDEL (Enzo Life Sciences). TMTC3 and TMTC4 cDNA was created by isolating RNA (HEK293T cells) and cloned into pcDNA3.1 A-, a plasmid harboring a C-terminal S-tag, using standard molecular biology techniques. All other chemicals were obtained from Sigma.

### Cell lines/Tissue Culture

HEK293T, 293A or COS7 cells were grown in DMEM supplemented with 10% fetal bovine serum, 100 U/mL penicillin and 100 mg/ml streptomycin. HEK293 cell lines (HEK293^SC^, HEK293^SC/POMT1,2^, and HEK293^SC/TMTC1,2,3,4^) were generously contributed by Dr. Henrik Clausen (University of Copenhagen) and grown in DMEM GlutaMax supplemented with 10% fetal bovine serum.

### In silico analysis of TMTC3 and TMTC4

The primary amino acid sequences of TMTC3 and TMTC4 were analyzed by UniProtKB and TPRpred to identify the number and position of putative TPR domains (Xu and Ng, 2015)(Karpenahalli *et al.*, 2007; UniProt: a worldwide hub of protein knowledge, 2019). Hydrophobic domains were identified by the ΔG software, which predicts transmembrane domains (Hessa *et al.*, 2007). Putative N-linked glycosylation sites were identified by NetNGlyc (Gupta, 2002).

### Affinity Purification and glycosylation assay

Transfected cells were lysed in MNT buffer (0.5% Triton X-100, 20 mM MES, 100 mM NaCl, 30 mM Tris-HCl [pH 7.5]). All steps were conducted at 4 °C. The post-nuclear supernatant (PNS) was isolated by centrifugation followed by pre-clearing with un-conjugated agarose beads for 1 hr. Cleared supernatant was incubated with S-protein agarose beads overnight and subsequently washed twice with wash buffer (0.05% Triton-X-100, 0.1% SDS, 300 mM NaCl, 100 mM Tris-HCl [pH 8.6]). After the final wash, glycosylation assays were performed by adding appropriate buffers and either mock, Endo H or PNGase F enzymes according to the manufacturer’s protocol. Finally, reducing sample buffer was added to all samples and they were analyzed by SDS-PAGE.

### Confocal Microscopy

Cells were fixed with 3.7% paraformaldehyde in phosphate buffered saline (PBS) for 15 min followed by permeabilization with 0.1% triton X-100 for 15 min at 25 °C. Slides were stained with the indicated primary antibodies followed by staining with appropriate Alexa Fluor 488 or 594 secondary antibodies in immunostaining buffer (10% fetal bovine serum in 1X PBS). Slides were rinsed and mounted onto cover slips with VectaShield (Vector Laboratories). Images were obtained with a Fluoview 1000 MPE, 1X81 motorized inverted research microscope (Olympus Inc.) equipped with a Hamamatsu C8484-05G camera. All images were acquired with a *Plan Apo N* 60x 1.42NA lens and processed by using the FV10-ASW and Adobe Photoshop software. For pan-cadherin staining, 24-well glass bottomed plates were prepared with 0.1% gelatin in PBS for 1 hr at 37 °C prior to coating each well with 10 μg/mL fibronectin (Sigma) overnight at 4 °C. Cells were allowed to adhere in the wells for 2 hr at 37 °C and subsequently fixed in methanol for 10 min at −20 °C. Cells were stained with the indicated primary antibodies followed by staining with appropriate Alexa Fluor 488 or 594 secondary antibodies in 1X PBS containing 0.1% Tween, 2% bovine serum albumin (Fisher) and 0.02% sodium azide. Cells were rinsed and then incubated with Hoescht (Abcam) for 1 hr at 25 °C. Images were obtained with a A1R: Nikon A1 resonant scanning confocal with TIRF module microscope equipped with an Andor Xyla camera. All images were acquired with a *Plan Apo IR* 60x 1.27WI lens and processed by using the Nikon Elements and Adobe Photoshop software.

### Alkaline extraction

Alkaline extraction was performed as previously described (Sunryd *et al.*, 2014). Briefly, radiolabeled cells were resuspended in ice-cold homogenization buffer (20 mM HEPES, 5 mM KCl, 120 mM NaCl, 1 mM EDTA, and 0.3 M sucrose [pH 7.5]) and passed through a 25-gauge needle 20 times. All subsequent steps were conducted at 4 °C. The homogenate was centrifuged at 1,000 g for 10 min to pellet the nuclear fraction. The remaining PNS was centrifuged at 45,000 rpm in Beckman rotor (TLA 120.2) for 10 min to separate the cytosol (supernatant) from the cellular membranes (pellet). The cellular membrane fraction was resuspended in homogenization buffer, and a portion of the resuspended membranes was incubated with 0.1 M Na_2_CO_3_ (pH 11.5) for 30 min on ice. The alkaline extracted portion was centrifuged at 65,000 rpm for 20 min through a sucrose cushion (50 mM triethanolamine, 0.3 M sucrose [pH 7.5]) to separate soluble proteins from membrane proteins in the supernatant and pellet, respectively. The pH was adjusted in the alkaline extracted sample with 1 M Tris-HCl (pH 7.5). An excess of MNT was added to all fractions, and immunoprecipitation or affinity precipitation was performed with protein-A sepharose and appropriate antisera or with S-protein agarose, respectively.

### Trypsin protection

Transfected cells were resuspended in cold homogenization buffer (10 mM HEPES [pH 7.4], 10 mM KCl, 1.5 mM MgCl, 5 mM sodium EDTA, 5 mM sodium EGTA and 0.25 M sucrose) and passed through a 25-gauge needle 20-times. All subsequent steps were conducted at 4 °C. The homogenate was centrifuged at 1,000g for 10 min to pellet the nuclear fraction. The remaining PNS was centrifuged at 45,000 rpm in Beckman rotor (TLA 120.2) for 10 min to separate the cytosol (supernatant) from the cellular membranes (microsomes). The microsomes were resuspended in homogenization buffer containing 0.1 M NaCl and 10 μg trypsin and/or triton X-100 was added to a final concentration of 0.1%. After incubation at 27 °C for 15 min, the reaction was quenched with 100 μg soybean trypsin inhibitor. Reducing sample buffer was added and analyzed via SDS-PAGE.

### Immunoblotting, endoglycosidase and affinity purification

Non-transfected and transfected cells were lysed in MNT and the PNS was isolated by centrifugation followed by total protein concentration determination using 595 nm Protein Assay Dye Reagent (Bio-Rad). All subsequent steps were conducted at 4 °C. Equal amounts of total protein were split into three fractions. Preclearing was performed on one of the fractions with control agarose beads for 1 hr prior to affinity purification with S-protein agarose beads overnight. The remaining two fractions were subjected to a glycosylation assay by adding appropriate buffers and either mock or PNGase F enzyme according to the manufacturer’s protocol. Following endoglycosidase treatment, affinity purification for glycoproteins using Con A (Sigma) was performed overnight. Beads were washed twice in wash buffer. Finally, reducing sample buffer was added to all samples and they were analyzed by SDS-PAGE. Proteins were transferred to a polyvinylidene difluoride membrane and immunoblotted with the appropriate antisera. Blots were developed and TIFF files were acquired using a LI-COR Odyssey CLx Imager. Densitometric quantification of western blots was performed using ImageJ software (Fiji). The amount of E-cadherin bound to Con A was calculated by dividing the amount of E-cadherin in the PNGaseF treated transfected cells with the amount of E-cadherin in the PNGaseF treated non-transfected cells (mock). The amount of E-cadherin in non-transfected cells was set to 100%. Error bars represent the standard deviation for three independent experiments. TMTC3 S-tag transfected samples were also subjected to a glycosylation and carbohydrate binding assay that included both PNGaseF and α1-2,3,6 mannosidase treatment. Briefly, transfected cells were lysed as described above and split into three fractions. Glycosylation assays were performed by adding appropriate buffers and either mock or PNGase F enzymes according to the manufacturer’s protocol and subsequent treatment with α1-2,3,6 mannosidase enzyme and appropriate buffers for 24 hr. Samples were analyzed as described above and the amount of E-cadherin bound to Con A was calculated by dividing the amount of E-cadherin in PNGaseF/α1-2,3,6-mannosidase treated samples by the amount of PNGaseF treated E-cadherin, which was set to 100%. Error bars represent the standard deviation for three independent experiments.

### qRT-PCR

HEK293A cells were treated with regular growth media or dithiothreitol (2 mM) for 2 hr or tunicamycin (1 μg/mL), thapsigargin (3 μM), brefeldin A (2.5 μg/mL) and MG132 (2.5 μM) for 24 hr prior to RNA isolation with RNAeasy Mini Kit (Qiagen). One μg of purified RNA was reverse transcribed into cDNA using the Protoscript II Reverse Transcriptase kit (New England Biolabs). Quantitative real time polymerase chain reactions (qRT-PCR) were performed in 20 μL reactions using the FastStart universal SYBR Green master (Rox) kit (Roche diagnostics Corp.) on an Mx3000P real-time PCR machine (Agilent Technologies Inc.) according to manufacturer’s instructions. Changes in mRNA levels were calculated using the change in cycle threshold value method with β-actin as the reference gene (Pfaffl, 2001). Statistical analysis of the data was calculated using GraphPad Prism 5.0 (GraphPad software) and significance between treatment groups was determined using unpaired T-tests.

The following primers were used: β-actin (5’ GCACTCTTCCAGCCTTCC 3’, 5’ TGTCCACGTCACACTTCATG 3’), TMTC1 (5’ GCTGTTTCTATTGGCCTTTCTC 3’, 5’ TGTCTCTTTCACCAGCATCG 3’), TMTC2 (5’ GATGTCTTTGTCTTTCACAGGC 3’, 5’ TGTTTCCCATCCAGTATAACCG 3’) TMTC3 (5’ TTTTCCTAAGCCATCCCCTG’, 5’ ACAAAACCACAAAAGAGGCTG 3’), TMTC4 (5’ CCCTCATTAAGTCCATCAGCG 3’, 5’ ATAACGAGAAATCCCAGGCC 3’), POMT1 (5’ GTGAACTACCTCCCGTTCTTC 3’, 5’ CACAGGGAGCAGAAGGATTT 3’), POMT2 (5’ GGCACTGGCCTATCAACTATC 3’, 5’ CAACAGATTCAGCCACCAAAC 3’) and BiP (5’ CTGCCATGGTTCTCACTAAAATG 3’, 5’ TTAGGCCAGCAATAGTTCCAG 3’).

### Cycloheximide chase analysis

HEK293T cells were transfected with S-tagged WT TMTC3 or mutant TMTC3 for 40 hr prior to incubation with 100 μg/mL of cycloheximide for indicated times. Cell lysates were prepared and the expression of S-tag TMTC3 was analyzed by Western blots. Briefly, transfected cells were lysed in MNT. Protein concentration of cell lysates was determined by using the 595 nm Protein Assay Dye Reagent (Bio-Rad). Western blots were carried out after 9% SDS-PAGE. After incubation with IRDye-conjugated secondary antibodies, the protein-antibody complexes were visualized by the LI-COR Odyssey CLx Imager and densitometric quantification of western blots was performed using ImageJ software (Fiji). The stability of TMTC3 was calculated by dividing the amount of TMTC3 WT or mutants in cyclohexmide treated samples (2-24 hr time points) by the amount of TMTC3 in untreated samples (0 time point), which was set to 100%. Error bars represent the standard error for three independent experiments.

### Cell Adhesion Assays

Substrates included bovine fibronectin (Sigma), E-cadherin (R&D Systems) and BSA (Fisher). Falcon Probind 96-well plates were coated with 25 and 50 μg/ml substrates in PBS overnight at 4□°C, rinsed with PBS, and blocked with 10 mg/ml BSA in PBS for 2 hr at 25 °C. Cells were harvested in serum free media by resuspension. 60,000 cells/well were seeded and the adhesion assay was performed for 1 hr at 37□°C in 5% CO_2_ in a humidified atmosphere. Non-adherent cells were washed away with serum free media. Adherent cells were fixed for 20 min with 4% formaldehyde in PBS, rinsed with PBS, and permeabilized with 0.1% triton X-100 for 20 min prior to staining with Hoescht (Abcam). Excess stain was removed with PBS. Each assay point was derived from three independent experiments.

### *Xenopus* embryo handling and media

Eggs were obtained from adult *Xenopus laevis*, fertilized, and cultured as described previously (Cousin *et al.*, 2008). Embryos were staged according to (Nieuwkoop and Faber 1994). Briefly, after in vitro fertilization, the eggs were dejellied in 2% cysteine (pH ∼8.3) and then cultured in 3% Ficoll in Modified Barth’s Solution (1X MBS) at 15 °C until they reached the desired stage for injection. Embryos were grown in 0.1X MBS at either 15 or 18° C. Time-lapse video microscopy was performed at 18° C.

### Microinjection experiments

Morpholinos (GeneTools, LLC) for Tmtc1, 2, 3 and 4 were designed to block translation of the respective proteins and contain a 3’-fluorescein modification by the manufacturer and injected (10 ng) at the one-cell stage. mRNAs of human TMTC3 WT and previously described mutants, cloned into pcDNA3.1 A-with a C-terminal S-tag, were transcribed using T7 polymerase following linearization of plasmids with EagI or DraIII. Transcripts were desalted on G-50 Nick columns (Pharmacia), extracted with phenol/chloroform, and ethanol precipitated. Transcripts were quantified by absorbance at 260 nm and resuspended at 0.2 μg/ml in DEPC-treated H_2_O. Transcripts (1 ng total) were injected at the one-cell stage, following injection of the TMTC3 morpholino.

### Embryo and cell imaging

Pictures of embryos were taken using the Zeiss Axiovert 200 M inverted microscope equipped with a Ludl xyz-stage control and a Hamamatsu Orca camera (Fig. 6) or BZ-X Analyzer equipped with Nikon 10X CFI PlanFluor lens and BZX710 camera (Keyence). Images were taken using either the AxioVision software (Zeiss) (Fig. 6) or the BZ-X capture software (BZ-X700 Analyzer)(Fig. 6). The diameter of the blastopore was measured using ImageJ software (Fiji) and quantified using GraphPad Prism 5.0 (GraphPad software). Statistical analysis between morpholino and TMTC3 construct injected groups was performed using one-way ANOVA and *, **, *** indicates a P-value of less than 0.05, 0.01 and 0.001, respectively. Error bars represent standard deviation.

### Protein extraction and analysis

Embryonic proteins were extracted from 10-20 frozen embryos using 1X MBS (Laskey *et al.*, 1977) containing 1% triton X-100, protease inhibitor cocktail (Sigma), 2 mM phenylmethylsulfonid fluoride, and 1 mM EDTA on ice followed by 16,000 g centrifugation for 30 min at 4 °C. Supernatant was either directly added to an equal volume of 2X Laemmli buffer containing 2% β-mercaptoethanol or precipitated with trichloroacetic acid (Sigma) and washed with acetone prior to resuspension in 2X Laemmli containing 2% of β-mercaptoethanol. Western blots were performed using standard techniques and the presence of translated TMTC3 constructs was assessed using anti-S-tag antisera.

## Acknowledgements

This work was supported by the National Institutes of Health under award number GM086874 to D. N. H.

## Conflicts of interest

The authors declare that they have no conflicts of interest with the contents of this article.

