## Supplemental Figures 1-7 for "ER transmembrane protein TMTC3 contributes to O-mannosylation of E-cadherin, Cellular Adherence and Embryonic Gastrulation"

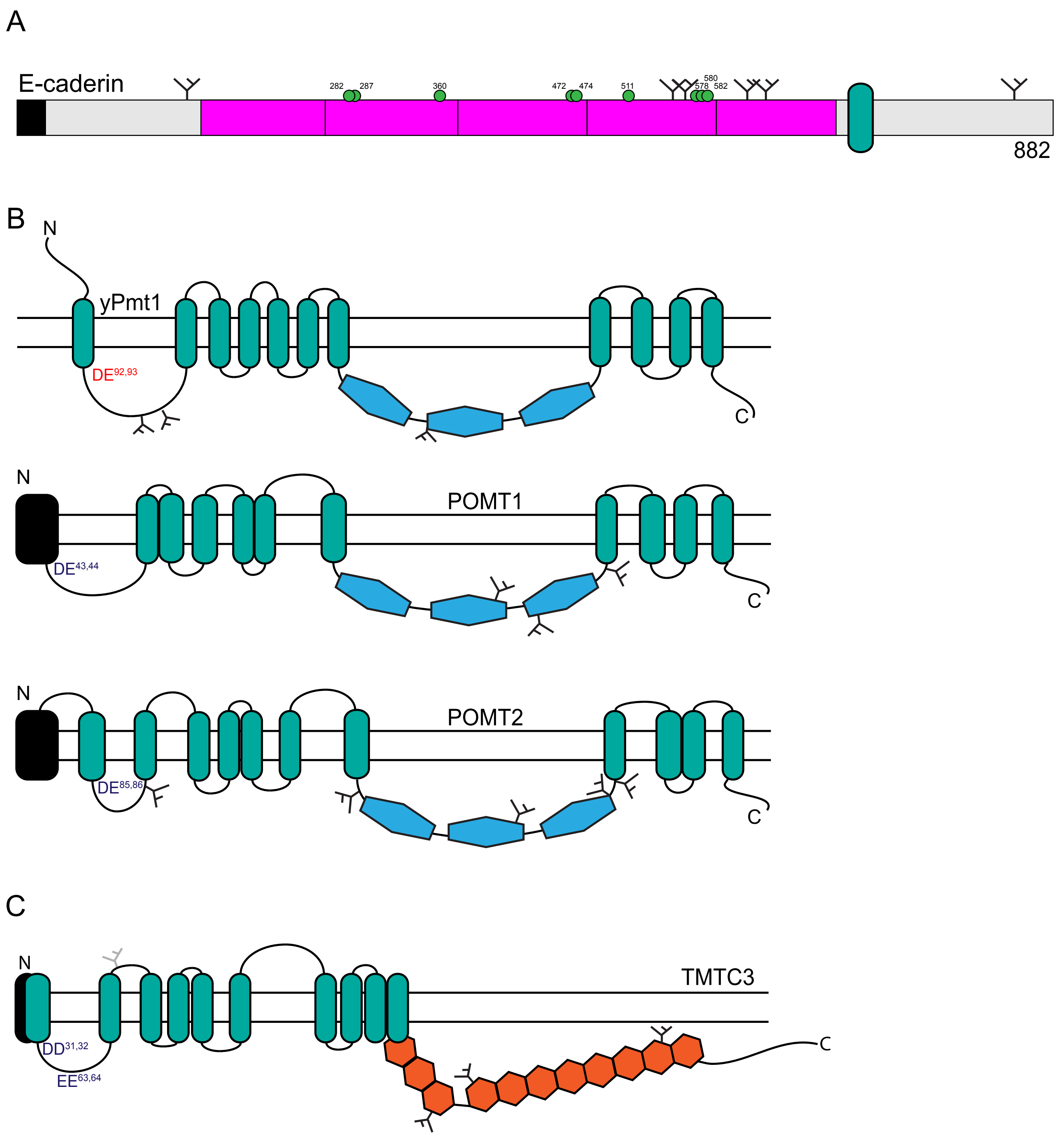


Supplemental Figure 1. *TMTCs and POMTs share similar functional domain architecture*. (A) The organization of E-cadherin with signal sequence (black), extracellular cadherin domains (pink) and transmembrane domain (teal) as designated. N-linked glycosylation sites are indicated by small black, branched structures and O-mannosylation sites by green circles. (B) The organization of yeast Pmt1 and predicted organization of human POMT1 and 2 with signal sequences (black), hydrophobic domains (teal) and MIR domains (blue) as designated. Predicted endogenous N-linked glycosylation sites are indicated by small black, branched structures and putative diacidic motif active sites are indicated in dark blue (red for active site that has been identified in yPmts).(C) Predicted organization of human TMTC3 with signal sequence (black), hydrophobic domains (teal), TPR domains (orange) and predicted active sites (diacidic DE and EE residues in dark blue) as designated.

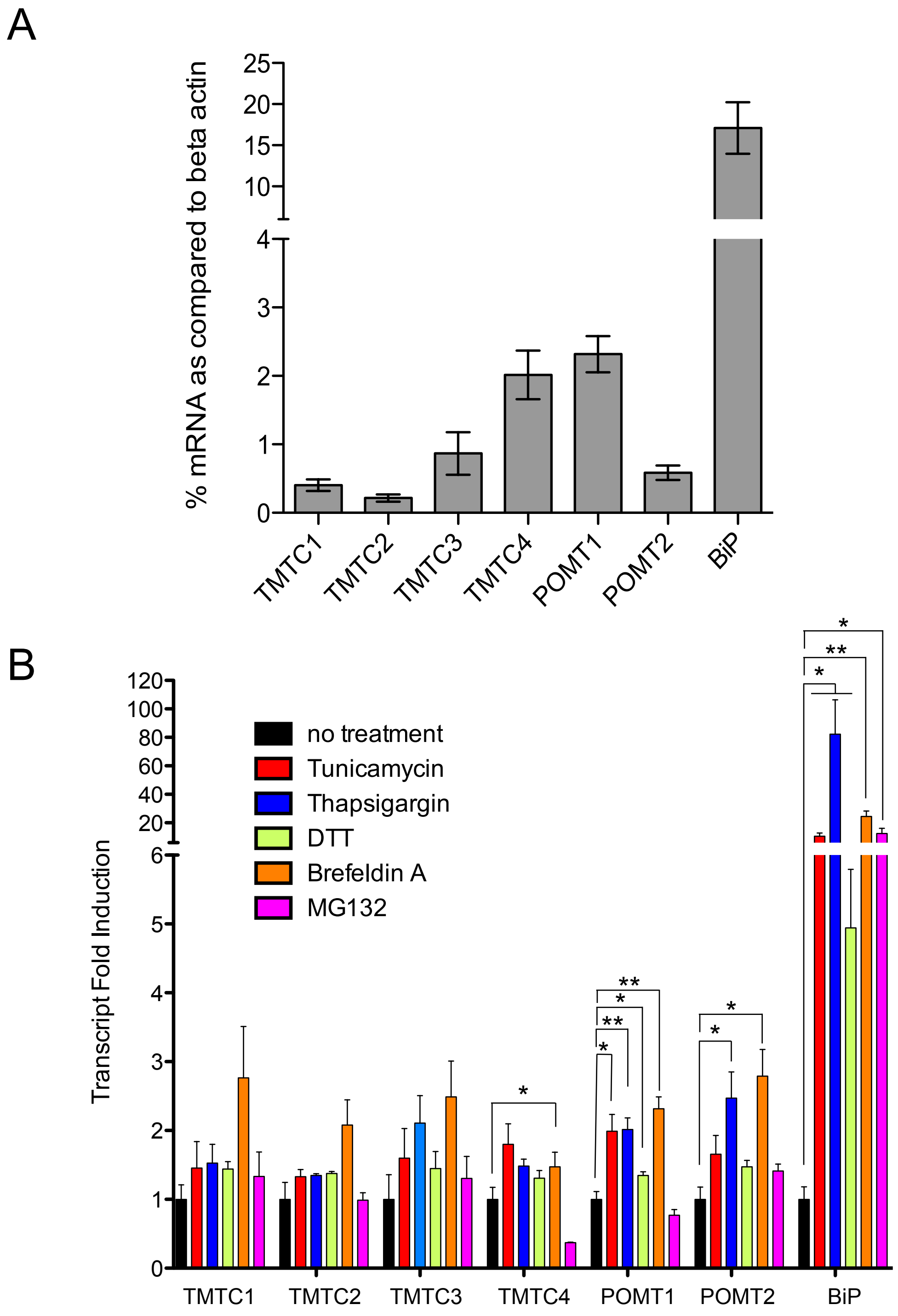


Supplemental Figure 2. *Basal transcript abundance and fold induction of TMTC proteins by ER stress.* (A) RNA from HEK293A cells, grown under normal conditions, was harvested. RNA was reverse transcribed to cDNA followed by qRT-PCR with appropriate primers. Basal mRNA abundance was assessed using β-actin as a reference. Error bars represent standard error from at least three independent experiments. (B) HEK293A cells were treated with regular growth media or with 2 mM DTT for 2 hr, 1 μg/mL tunicamycin, 3 μM thapsigargin, 2.5 μg/mL brefeldin A or 2.5 μM MG132 for 24 hr prior to RNA purification. RNA was reverse transcribed to cDNA followed by qRT-PCR with appropriate primers, and changes in gene expression were calculated using β-actin as a reference. Statistical significance between treatment groups was determined using an unpaired T test. * and ** indicates a P-value of less than 0.05 and 0.01, respectively. Error bars represent standard deviation from at least three independent experiments.

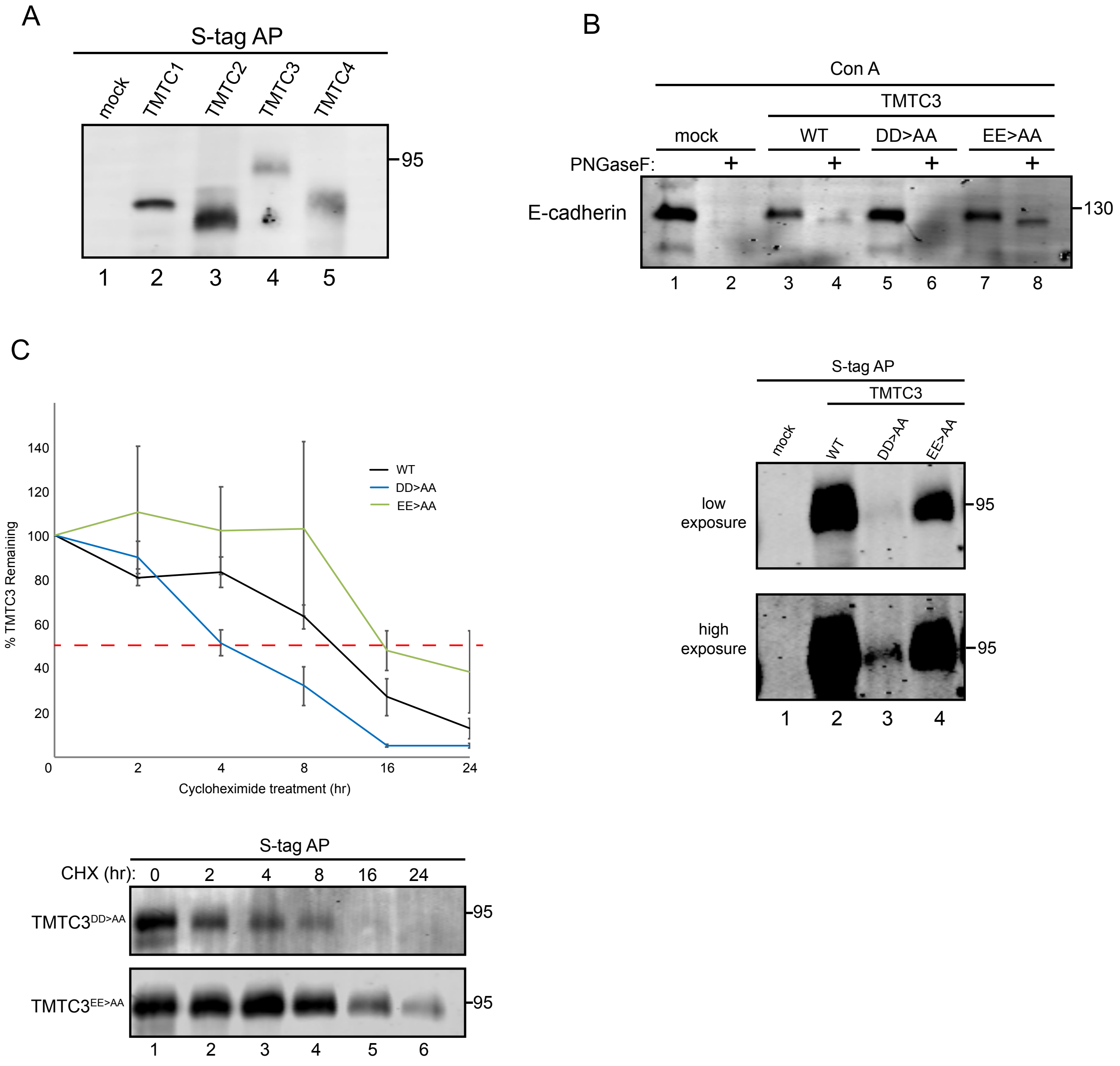


Supplemental Figure 3. *Deletion of TMTCs and E-cadherin*. (A) S-tagged TMTC1, 2, 3 or 4 cDNA was transfected into HEK293[SC/TMTC1,2,3,4] cells. Cell lysates were collected and one third was subjected to affinity purification with S-protein agarose. Samples were then analyzed by 6% SDS-PAGE and immunoblotted for the S-tag epitope to assess levels of TMTC1, 2, 3 and 4, respectively. (B) HEK293[SC/TMTC1,2,3,4] cells were transfected with S-tagged TMTC3 WT or putative active site mutant (DD>AA or EE>AA) cDNA. Lysates were collected and split, one third subjected to treatment with endoglycosidase PNGaseF (lanes 2, 4, 6, 8 and 10), prior to pulling down glycosylated proteins with concanavalin A (Con A). One third was subjected to affinity purification with S-protein agarose. Samples were then analyzed by 6% SDS-PAGE and immunoblotted with E-cadherin antisera (top blot) or S-tag antisera to assess levels of TMTC3 (lower blots). Lower panel represents the upper panel S-tag blot at a higher exposure to see the diminished expression of the DD>AA mutant (lane 3) (C) HEK293T cells were transfected with S-tagged TMTC3 WT or putative active site mutant cDNA as indicated. Cells were treated with 100 µg/mL cycloheximide for the indicated time prior to collection in lysis buffer. Samples were subjected to affinity purification with S-protein agarose and analyzed via 9% SDS-PAGE and immunoblotted for the S-tag epitope. Time points were collected at 0, 2, 4, 8, 16 and 24 hr after cycloheximide treatment. The amount of TMTC3 protein remaining at 2, 4, 8, 16 and 24 hr was quantified and normalized to the starting material (0 h) and averaged from three independent experiments. Error bars represent standard error of the mean (upper panel).

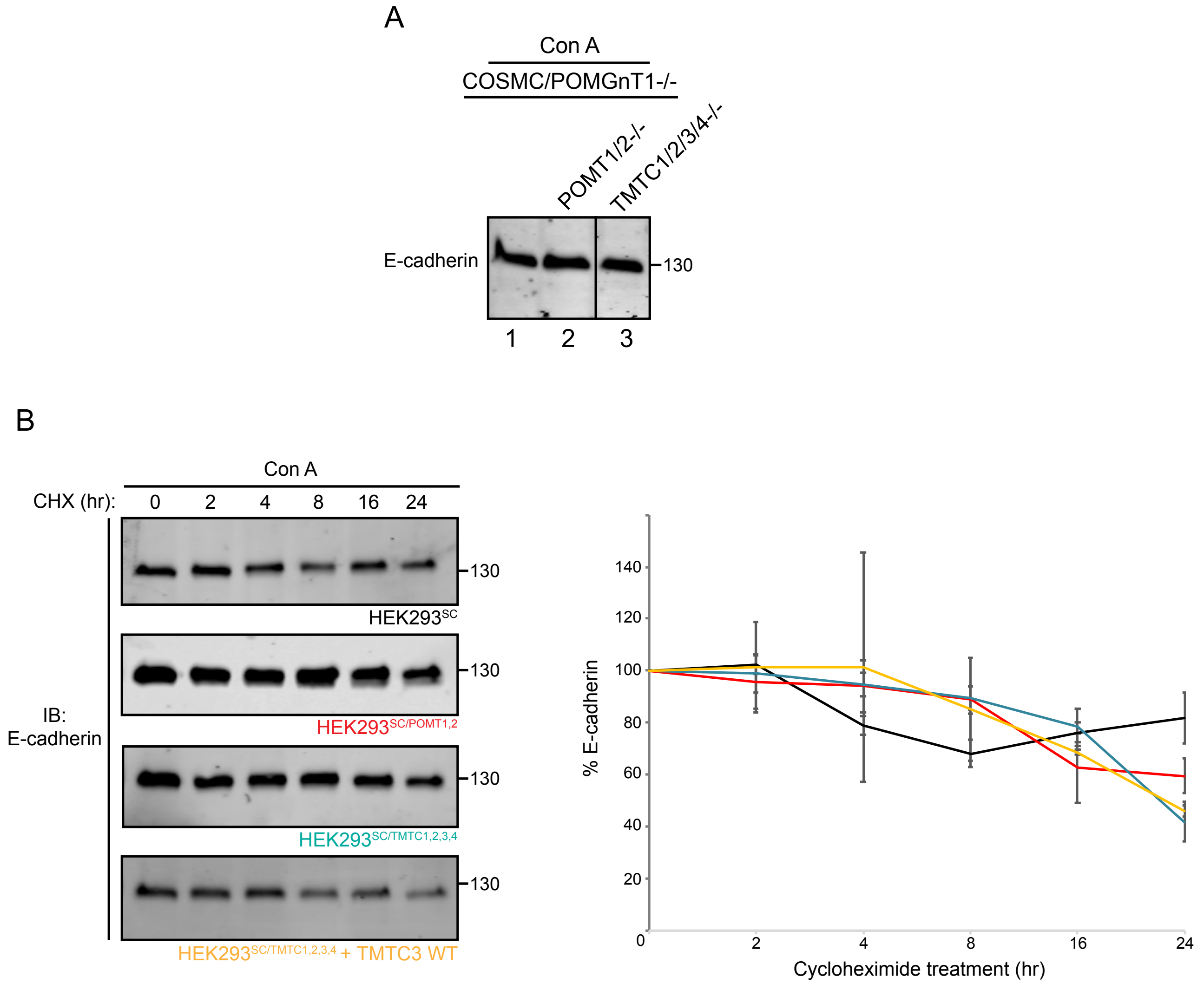


Supplemental Figure 4. *Deletion of TMTCs and E-cadherin*. (A) Expression and stability of E-cadherin was assessed in O-mannosyltransferase knockout cells to determine effects of O-glycosylation. HEK293$^{SC}$, HEK293$^{SC/POMT1,2}$ and HEK293$^{SC/TMTC1,2,3,4}$ cells were lysed and total protein amount was assessed via Bradford assay. Equal amounts of protein were subjected to pull-down via Con A prior to samples being analyzed by 9% SDS-PAGE and immunoblotted with E-cadherin antisera to monitor levels of E-cadherin. (B) Previously described cell lines were treated with 100 μg/mL cycloheximide for indicated time (0, 2, 4, 8, 16 and 24 hr) prior to lysis, total protein quantification and subsequent affinity purification with Con A. Samples were then analyzed by 9% SDS-PAGE (left panel) and the amount of E-cadherin remaining at 2, 4, 8, 16 and 24 hr was quantified and normalized to the starting material (0 h) (right panel). Error bars represent standard error of the mean.

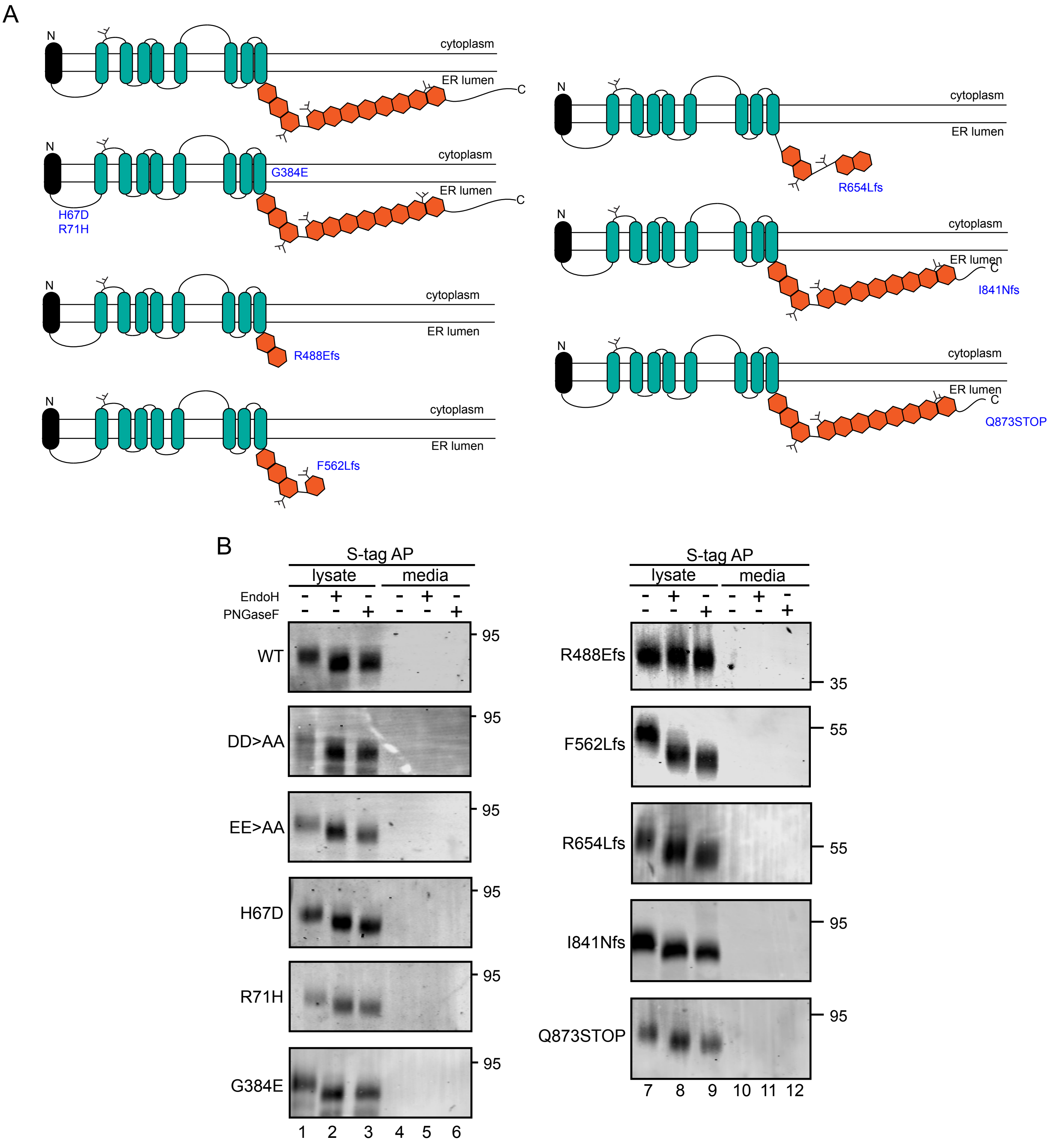


Supplemental Figure 5. *TMTC3 disease variant predictive outcomes and carbohydrate analysis*. (A) The organization of TMTC3 WT and disease variants signal sequence (black), hydrophobic domains (teal), TPR domains (orange). N-linked glycosylation sites are indicated by small black, branched structures. Mutations are indicated in blue. (B) HEK293T cells were transfected with S-tagged TMTC3 WT and disease variants cDNA as indicated and were affinity purified from the media and the lysed cells using S-protein agarose. Samples were then subjected to a glycosylation assay with either EndoH (lanes 2, 5, 8 and 11) or PNGaseF (lanes 3, 6, 9 and 12) digestion as indicated. Reducing sample buffer was added, and the samples were analyzed by a 6% SDS-PAGE.

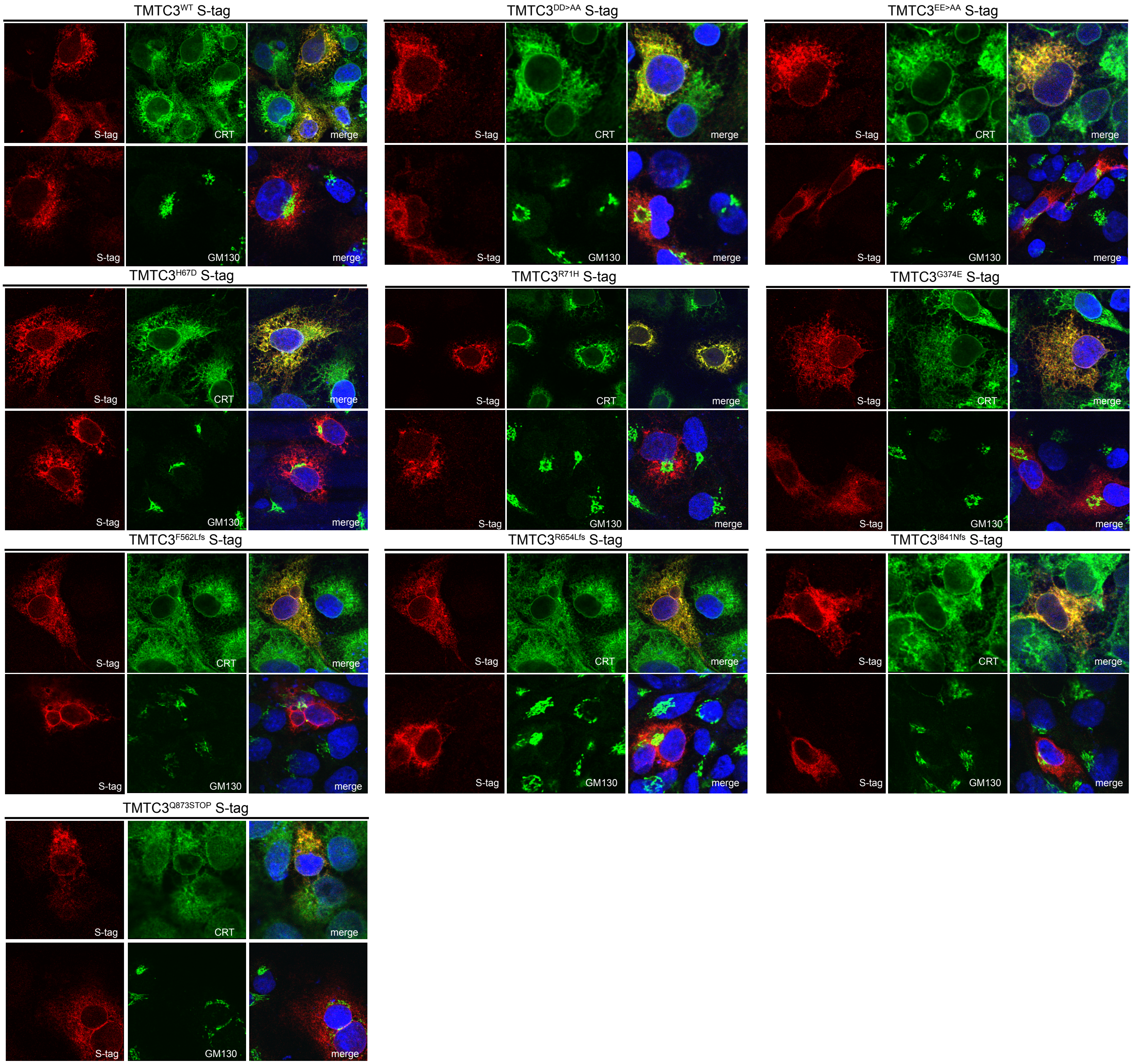


Supplemental Figure 6. *Cellular localization of TMTC3 disease variants*. Cellular localization of TMTC3 disease variants was investigated by confocal microscopy. Cos7 cells were transfected with TMTC3 disease variant cDNA. Fixed cells were stained with S-tag, ERp57 (ER) or GM130 (Golgi) antisera. Nuclei were visualized by DAPI staining (blue).

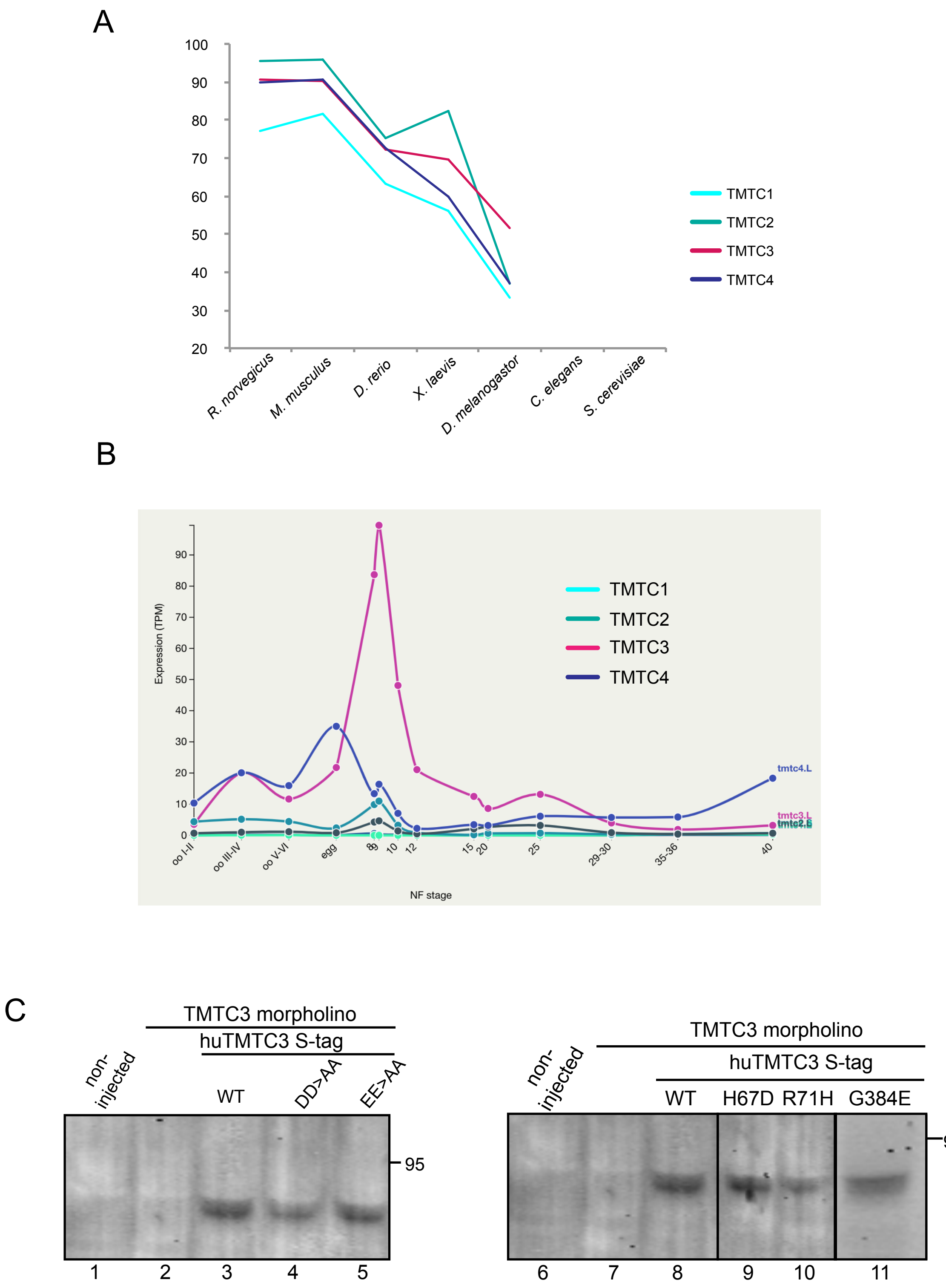


Supplemental Figure 7. *TMTC conservation in commonly studied species and RNA expression in Xenopus laevis*. (A) Sequence identity (%) of the TMTC proteins in commonly studied species was assessed by comparing amino acid sequences of each species listed to the human sequence. (B) RNA expression of *Tmtc1-4* during *Xenopus laevis* development up to stage 40. These expression profiles were based on data collected in the study by Session *et. al* in Nature 2016. (C) Embryos were injected with Tmtc3 morpholino at the one cell stage approximately 45 min after fertilization. Selected embryos were subsequently injected with human TMTC3 RNA (lanes 3, 4, 5, 8, 9, 10 and 11). Non-injected and indicated injected embryos were collected at stage 12 and resuspended in Modified Barth's Saline (MBS) buffer with 1% triton X-100. Total protein was precipitated with 10% trichloroacidic acid, resuspended in reducing sample buffer, analyzed by a 9% SDS-PAGE and immunobloted for the S-tag epitope.
